# Thermal variation influences the transcriptome of the major malaria vector *Anopheles stephensi*

**DOI:** 10.1101/2024.05.27.596085

**Authors:** Ashutosh K Pathak, Shannon Quek, Ritu Sharma, Justine C Shiau, Matthew B. Thomas, Grant L. Hughes, Courtney C Murdock

## Abstract

The abundance and distribution of ectotherms is being shaped at unprecedented scales by human-induced climate change. Unraveling molecular adaptations to thermal variation is prescient. Vector-borne diseases are of particular concern and although mosquitoes are responsible for most of the 621 million cases annually, their response to temperature is not well characterized. We describe the transcriptome of the major vector for human malaria, *Anopheles stephensi* maintained under a diurnal temperature regime (DTR) totaling 9°C and daily means of 20°C, 24°C and 28°C respectively. Gene expression profiles were examined in the midguts and carcasses of adult mosquitoes from one to 19 days post-blood meal. Differences in temporal expression profiles at each temperature revealed a total of 3,106 and 3,590 genes in the carcasses and midguts respectively and analyzed further to make three inferences. First, we identified genes with shared temporal expression profiles across all three temperatures, suggesting their indispensability to mosquito life history. Second, tolerance to 20 DTR 9°C and 28 DTR 9°C was associated with a larger and more diverse repertoire of gene products compared to 24 DTR 9°C; assuming physiological costs scale accordingly, this finding could explain the fitness trade-offs underlying the unimodal effect of temperature. Third, the long duration of our study revealed two general trends in gene expression, representative of a coordinated, tissue- and temperature-specific response to blood meal digestion, managing oxidative stress, and reproduction. Our results suggest how temperature could regulate the mosquito’s capacity to transmit *Plasmodium* parasites and offer a reference point for understanding ectotherm adaptation.

## INTRODUCTION

Climate change is expected to lead to the redistribution of mosquito species on a global scale. While climate change is the cumulative effect of several environmental factors, changes in temperature have been demonstrated in numerous studies to be one of the most important factors constraining the distribution and abundance of ectothermic organisms (Brown, Pascual, Wimberly, Johnson, & Murdock, 2023; J. L. Chen & Lewis, 2024; Deutsch et al., 2008; González-Tokman et al., 2020; Harvey et al., 2023; Kingsolver & Buckley, 2017; Sunday, Bates, & Dulvy, 2012). For mosquitoes, and ectotherms more generally, the relationship between temperature and the performance of traits relevant for lifetime fitness is unimodal, where performance increases until the optimum temperature is reached, after which it starts to decline (reviewed by (Mordecai et al., 2019)). Based on the metabolic theory of ecology, as temperatures warm toward the thermal optimum, the rate and efficiency of enzymatic processes will increase. Beyond the thermal optimum, enzymatic processes are predicted to become less efficient due to protein misfolding or desiccation stress, until the critical upper temperature is reached where organismal death occurs (Brown et al., 2023). However, the physiological mechanisms underlying this unimodal response remain unclear. Identifying these mechanisms is pertinent for at least three reasons. First, to understand the mechanisms underpinning plasticity of mosquitoes to respond to different temperatures. Second, to provide more reference points for local adaptation within and between species. Third, to determine if the gene(s) that ultimately mediate these physiological responses could be selectively inherited in response to the current trends in climate change (Logan & Cox, 2020).

Despite the implications for public health, few studies have identified the genetic basis for how mosquitoes respond to temperature (Ferreira et al., 2020; Wimalasiri-Yapa et al., 2021). Several species within the genus *Anopheles* vector parasitic, unicellular eukaryotes of the genus *Plasmodium*, the etiological agent of malaria in several vertebrate species, including humans where it continues to cause significant morbidity and mortality (WHO, 2021). Environmental temperature regulates the rates of parasite establishment and development, as well as mosquito traits that govern mosquito population and transmission dynamics (Miazgowicz et al., 2020; Murdock, Blanford, Luckhart, & Thomas, 2014; Murdock, Sternberg, & Thomas, 2016; Pathak, Shiau, Thomas, & Murdock, 2019; Shapiro, Whitehead, & Thomas, 2017; Villena, Ryan, Murdock, & Johnson, 2022). The thermal performance curves of mosquitoes are also unimodal with mosquito species differing significantly in thermal breadth (Johnson et al., 2015; Kirk, O’Connor, & Mordecai, 2022; Miazgowicz et al., 2020; Shapiro et al., 2017; Villena et al., 2022). Amongst *Anopheles*, *An. stephensi* is considered a major vector for *Plasmodium* parasites (Sinka et al., 2020) in South Asia and more recently, urban centers of Africa (Sinka et al., 2020; Tadesse et al., 2021). In addition to this species’ relatively unique ability to develop in water storage containers, its thermal breadth is also thought to have contributed to its success in establishing in warmer and drier environments than other *Anopheles* (Sinka et al., 2020). Whether this thermotolerance associated with *An. stephensi* can be explained by transcriptional responses has not been assessed.

Physiological mechanisms underlying fitness traits are often the product of multiple genes and interacting pathways (Oomen & Hutchings, 2022; Todd, Black, & Gemmell, 2016). However, most of our understanding of the thermal responses of mosquitoes is derived from species housed at constant temperatures. For the same mean daily mean temperatures (e.g., 24°C), a few studies have shown the total number of genes and overall profiles can vary significantly between ectotherms housed at constant temperatures as compared to thermally fluctuating conditions, across a wide range of mean daily temperatures (15-31°C), with daily variations ranging from 3°C to 8°C (Breitenbach, Bowden, & Paitz, 2022; Salachan & Sorensen, 2022; Sorensen, Schou, Kristensen, & Loeschcke, 2016). Resolving responses in realistic environments is essential to the understanding of ectotherm adaptations.

In the current study, we use a bulk RNAseq approach to describe how variation in temperature modulates global gene expression profiles in *An. stephensi* on a spatial and temporal basis. Mosquitoes were maintained under field-relevant diurnally fluctuating temperature regimes around mean temperatures of 20°C, 24°C and 28°C, with 24°C being closest to the predicted thermal optimum (*T_opt_*) for malaria transmission potential (∼25°C) (Pathak et al., 2019; Villena et al., 2022). Patterns of global gene expression were determined from midguts and carcasses collected at various points from 1 to 19 days post-blood meal (Figure 1). This study demonstrates several important insights. Overall gene expression at the cool and warm temperatures reflected larger and more diverse transcriptional regulation. We also observe spatial regulation of genes in a tissue specific manner in response to temperature variation, and that temperature variation significantly impacted the transcriptomic response to blood meal digestion with important implications for mosquito egg production, gonotrophic cycle length, generation time, and mosquito population dynamics. Finally, our results suggest transcriptomic responses to temperature variation under diurnally fluctuating environments may be distinct from those characterized under constant temperature variation, with implications for our understanding of phenotypic plasticity to temperature variation in the field.

**Figure 1:**
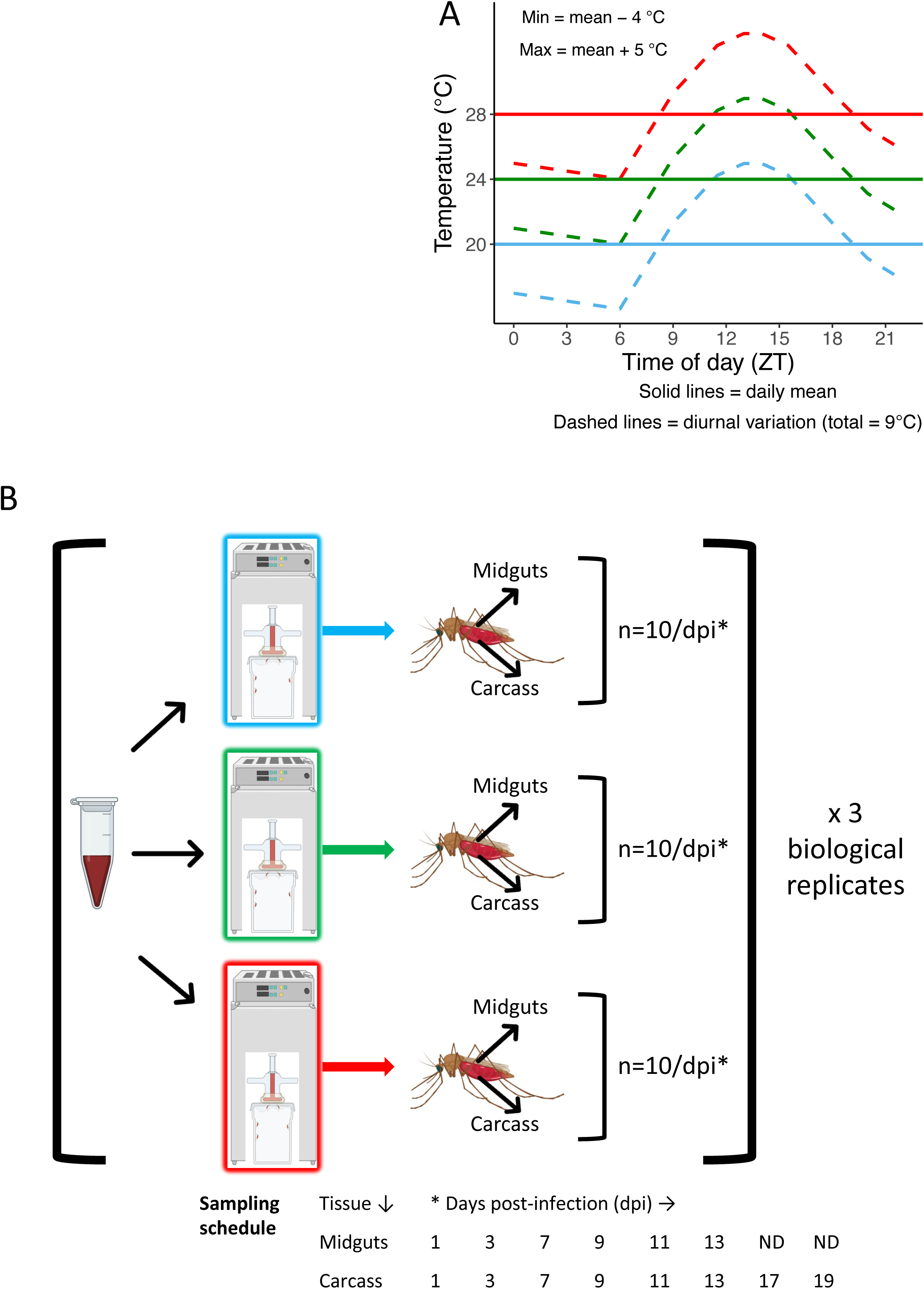
(A) Schematic of temperature regime used in this study and (B), overview of study design partly created with BioRender.com.

## METHODS

### Mosquito husbandry, blood feeds and sample collections

For schematic of study design, refer to Figure 1. Mosquito husbandry and blood feeding was performed as described previously (Pathak, Shiau, Thomas, & Murdock, 2018; Pathak et al., 2019). Briefly, 500, 3-5-day old female, host-seeking *An. stephensi* were aspirated into each of three polyester cages (32.5cm^3^ BugDorm, Taiwan). Mosquitoes were acclimated to the three temperatures for 48 hours prior to the blood meals. Blood feeds were performed with washed human RBCs reconstituted with human serum (Valley Biomedical, Winchester, VA), as described previously (Pathak et al., 2018, 2019). Mosquitoes were then returned to the respective temperatures. At day 1, 3, 7, 9, and 13 post-blood meal, midguts were dissected from 10 individuals and pooled into 50μL of RNAlater (Thermo-Fisher Scientific, Waltham, MA). After dissecting out midguts, the remaining mosquito tissue (referred to as carcasses herein) from these 10 individuals was pooled into 0.5ml of RNALater; note that carcasses were also collected on days 17 and 19 post-blood meal. After overnight storage at 2-4°C, samples were transferred to -80°C until RNA extraction. These procedures were repeated over three biological replicates. Each replicate provided a total of 36 samples over the various time points (5 midgut and 7 carcass pools at each temperature per replicate). The three replicates together provided 108 samples for RNAseq.

### RNA purification and sequencing

Total RNA was purified from the midguts and carcasses with commercially available kits (PureLink RNA Mini Kit) as recommended by the manufacturer (Thermo-Fisher Scientific, Waltham, MA). RNA sequencing (RNAseq herein) was performed by the Georgia Genomics and Bioinformatics Core (GGBC) at the University of Georgia (Athens, GA). Concentration and integrity of total RNA was assessed with a Bioanalyzer (Thermo-Fisher Scientific, Waltham, MA). Libraries for RNAseq were prepared for each sample with a KAPA stranded mRNA-seq kit as per manufacturer’s recommendations (Roche, Indianapolis, IN); briefly, mRNA was first selected with oligo-dT beads, followed by fragmentation of the RNA prior to generating cDNA using random hexamer priming. Quality and quantity of each library was estimated with a qubit or plate reader before pooling for RNAseq. Pooled libraries were run on a NextSeq 550 (Illumina Inc., San Diego, CA). De-multiplexing and trimming of adapter and barcode sequences were performed on BaseSpace (Illumina Inc., San Diego, CA).

### Data analysis

Sequence data were obtained in fastq format, and read quality was confirmed using FASTQC v0.11.5 (Andrews, 2010), with all samples showing an average PHRED score >30 and no adapter sequences detected. Samples that showed < 1 million sequenced reads were not included in subsequent differential expression analysis due to concerns about accurate quantification of gene expression. Sequences were aligned to the *Anopheles stephensi* “Indian” strain reference genome (genome version GCA_000300775.2, annotation version AsteI2.3) using the program Hisat2 (v2.1.0) (Kim, Paggi, Park, Bennett, & Salzberg, 2019). FeatureCounts v2.0.1 (Y. Liao, Smyth, & Shi, 2014) was used to obtain raw read count data from the sequencing files. The resulting feature count table was then processed using the program DESeq2 (v1.30.1) (Love, Huber, & Anders, 2014) in R (v4.3.0) (R Core Team, 2021). DESeq2 automatically performs filtering of genes with low read counts to increase statistical power, as well as normalization of gene expression based on the raw count data. This was used to generate a PCA plot and sample-to-sample distance matrixes in DESeq2 for further quality control and identification of sample clustering. Five samples appeared as distinct outliers in both analyses (no clustering with any biological replicate or any other sample) and were thus excluded from subsequent differential expression analysis. Despite sample removals based on low read counts or outliers from the remaining samples, all experimental conditions had a minimum of two biological replicates for differential expression analysis.

To investigate the impact of temperature on gene expression in mosquitoes over the course of the experiment (19 days), we utilized the DESeq2’s in-built Likelihood Ratio Test (LRT) (Love et al., 2014). This type of test can be useful in analyzing time-course experiments, as it evaluates expression change across multiple experimental factors. The LRT test looks at gene expression profiles and identifies genes that show differences in both expression as well as patterns across temperatures. I.e., genes that show changes in expression level, but have the same broad pattern over time, would be considered as non-significant.

To first investigate gene changes over time at the three different temperatures, we treated each mosquito colony at the different temperatures as a separate time-series experiment, with design formula of ∼Time, reduced formula of ∼1, and adjusted p-value cutoff of 0.05. No thresholds of log_2_ fold-changes were applied, as we were also interested in investigating the influence of small changes across an entire pathway. This analysis was performed for both guts and carcasses separately. The resultant list of differentially expressed genes was then used to create an Upset plot using the UpsetR package v1.4.0 (Conway, Lex, & Gehlenborg, 2017) to visualize the number of differentially expressed genes at the different temperatures and overlaps among them. These overlaps indicated the presence of clusters of genes that may show differential expression based on temperature.

To identify genes that may be clustering together based on gene expression, as well as visualize these changes over time, we utilized a technique known as K-means clustering via the MFuzz package (v2.60.0) (L. Kumar & M, 2007). This is a technique used for soft-clustering analysis of data and allows subsequent illustration of gene expression profiles (GEPs) in time-series data. MFuzz requires as input normalized count-data, taken from DESeq2 (Love et al., 2014), as well as a user-defined number of ‘cluster cores’, which defines the number of GEPs to be drawn. Too few cluster cores will result in reduced sensitivity for finding unique GEPs, while too many cluster cores will result in redundant patterns. Normalized count data for the statistically significantly differentially expressed genes for each of the different temperatures and tissues were used- other genes that did not meet an adjusted p-value of 0.05 were discarded for this analysis. For the identification of cluster cores, we performed clustering analysis using an increasing number of cluster cores, and scored the correlation of the resulting clusters using the ‘cor’ function of base R stats package (R Core Team, 2021). The number of cluster cores to use was determined by a maximum correlation score of 0.85, where a correlation score of 1 indicates perfect match of the gene expression profile, and a score of -1 indicates a perfect inversion of the profile. This resulted in between 3 to 5 cluster cores depending on the analysis being investigated, with their corresponding expression profiles drawn using MFuzz (L. Kumar & M, 2007) and the ggplot2 package (Wickham, 2016). For identifying changes in expression, identified GEPs were compared against one another, both in terms of profile and the gene lists associated with them to see how genes may change expression profiles depending on mosquito rearing temperature. Gene lists from each GEP were then processed for gene ontology enrichment to assign function for each profile using the g:Profiler web server (Reimand, Kull, Peterson, Hansen, & Vilo, 2007), with a significance cutoff of 0.05.

## RESULTS AND DISCUSSION

From all experimental conditions, we acquired between 4,302 to 128,645,661 reads, with five samples showing less than one million sequenced reads. Furthermore, initial PCA clustering of counts results showed six samples that appeared as clear outliers in the data and did not cluster with any other data point. These eleven samples were removed from downstream differential expression analysis.

### Variation in transcriptomes is driven by the blood meal and tissue-specific expression profiles

The effect of temperature on overall gene expression profiles at each time point in carcasses and midguts was investigated. The PCA plot indicated two major clustering factors that together accounted for 61% of the overall variation in the data (Figure 2A). The first principal component (PC1) was site of sampling (36% variance), with gene expression profiles at the site of blood meal ingestion (midguts) distinct from the systemic responses (carcasses) (Figure 2, ovals with solid line); in general, profiles were more variable between the midguts. The second principal component was time (25% variance), with the blood meal eliciting distinct gene expression profiles at day 1 compared to the later time points (Figure 2A, ovals with dotted line). While ingestion of a blood meal has a large effect on the transcriptomes of several mosquito species, tissue specific effects have also been reported previously (Domingos, Pinheiro-Silva, Couto, do Rosario, & de la Fuente, 2017; Hixson et al., 2022; Maccallum, Redmond, & Christophides, 2011).

**Figure 2:**
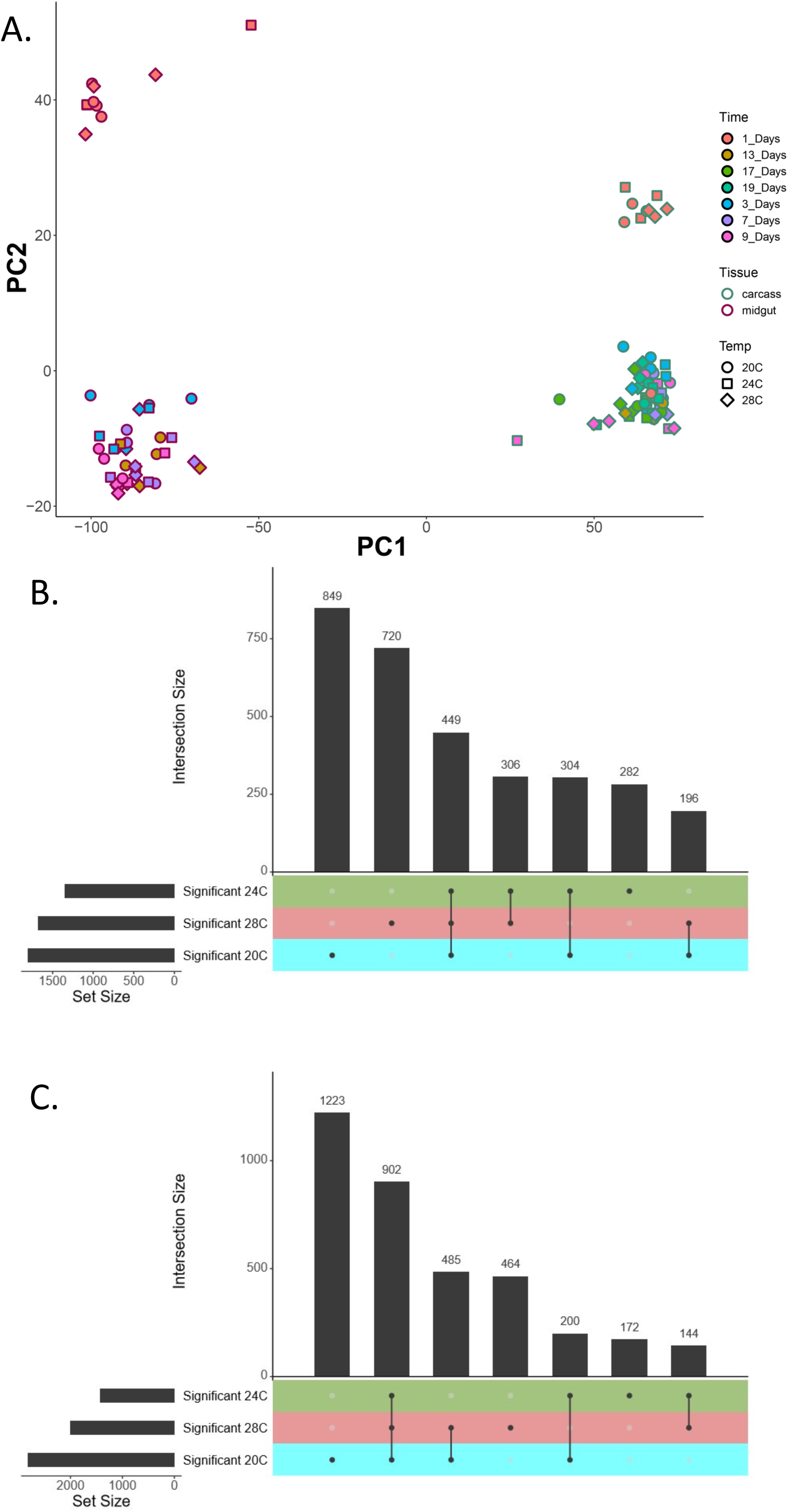
Summary plots showing characteristics of differentially expressed genes in carcasses and midguts of *An. stephensi* across different temperatures and times. (1A): PCA plot illustrating similarities and differences between samples. Note the distinction between carcasses and midguts (solid oval outline), as well as Day 1 versus other time-points (dashed oval outline). (1B): UpsetR plot showing how genes in carcasses show unique or shared statistically significant differential expression at certain temperatures between 20 DTR 9°C, 24 DTR 9°C and 28 DTR 9°C. UpsetR plots are an alternative method of illustrating complex Venn diagrams-each of the colored rows indicates a separate ‘set’, horizontal bar chart indicates set size, ball-and-stick diagram represent the different intersections, and vertical bar chart indicates intersect size. In this case, note how both 20 DTR 9°C and 28 DTR 9°C sets have the highest number of uniquely differentially expressed genes (849 and 720 genes respectively), followed by the intersect of all three temperature sets, with 449 genes shared between all temperatures. (1C): UpsetR plot showing the distribution of genes in midguts. Note the difference with carcasses- the largest set size comes from genes uniquely expressed at 20 DTR 9°C (1223 genes), followed by genes that are shared between all three temperatures (902 genes), followed by genes shared between 20 DTR 9°C and 28 DTR 9°C (485 genes).

### Tolerance to cooler (20 DTR 9°C) and warmer (28 DTR 9°C) temperatures was associated with a larger repertoire of gene products than the more optimal 24 DTR 9°C

Likelihood Ratio Tests was used to analyze the data as a time-series to capture genes that showed statistically significant deviation in expression patterns at the three different temperatures. This also allowed for subsequent clustering of these patterns and comparisons between the different temperatures analyzed. From this analysis, 1,798, 1,341 and 1,671 genes were identified as statistically significantly differentially expressed in the carcasses at 20, 24 and 28 DTR 9°C respectively. In the midguts, 2810, 1418 and 1995 genes were differentially expressed at 20, 24 and 28 DTR 9°C. These genes were not necessarily unique per temperature and organ assayed, with up to 902 genes shared across the analyzed temperatures in various combinations (Figure 2B, 2C1).

Next, LRTs were used to identify genes that show statistically significant changes in expression patterns over time at the three different temperature treatments. In the carcasses, a total of 3,106 unique genes were identified across the different temperatures, when aggregated over time. Of the genes unique to each temperature, a significantly higher number of differentially expressed genes were recovered at the low and high temperatures of 20 DTR 9°C and 28 DTR 9°C (849 and 720 respectively), compared to 24 DTR 9°C (282) (Figure 2B). While 449 genes were shared across all three temperatures, the remaining genes showed similar patterns between at least two of the three temperatures (e.g., the 306 between 24 DTR 9°C and 28 DTR 9°C, or the 304 genes across 20 DTR 9°C and 24 DTR 9°C) (Figure 2B).

Analyses of the midguts identified a total of 3,590 unique genes with clear differences in expression levels over the three temperatures aggregated over time. Unlike the carcasses, the largest group of significant genes were found at 20°C DTR 9°C (1,223 genes), with the second largest group represented by genes that were shared across all temperatures (902 genes) (Figure 2C). While 464 genes were still unique to 28 DTR 9°C, like the carcasses, only 172 genes were expressed exclusively in the midguts of mosquitoes housed at 24 DTR 9°C. The remaining 829 genes from the 3,590 total were shared between at least two of the three temperatures (Figure 2C). Transcriptomics of larvae of the major African vector *An. gambiae* showed that, compared to the 2L+^a^ variants, the higher resistance to thermal stress in populations with the chromosomal inversion 2La was associated with expressing a larger repertoire of differentially expressed genes (Cassone et al., 2011). Future research determining the extent to which this larger repertoire exerts a downstream effect on mosquito physiology or places greater demands for resources may prove to be useful. For instance, it can suggest pathways and mechanisms underlying the reduced fitness at sub-optimal temperatures, and the unimodal effect of temperature for mosquitoes, and ectotherms in general.

### Temporal classification of differentially expressed genes reveals clusters of shared and unique genes in the carcasses and midguts of mosquitoes at all three temperatures

To get a higher resolution of how temperature affects variation in the temporal dynamics of gene expression, specific group(s) of unique and shared genes identified above (Figure 2) were analyzed using K-means clustering. This separated each group of genes into 3-5 ‘clusters’ based on the similarity in expression profiles over time. We first looked at the temporal clustering profiles of the genes uniquely expressed in carcasses at each temperature. The 849 (20 DTR 9°C), 282 (24 DTR 9°C), and 720 (28 DTR 9°C) genes uniquely expressed in the carcasses at each temperature (Figure 2B) could be separated into 3-4 clusters per temperature (Figure 3A-C), with between 64 to 347 genes in each cluster (Table 1). Similar clusters were also derived from the differentially expressed genes in the midguts of mosquitoes at days 1, 3, 7, 9 and 13 after the blood meal. For the 1,223 (20 DTR 9°C), 172 (24 DTR 9°C) and 464 (28 DTR 9°C) genes uniquely expressed in the midguts at each temperature (Figure 2C), 3-5 gene clusters were identified per temperature (Figure 3D-F) with sizes ranging from 42-313 genes per cluster (Table 1). For both carcasses and midgut gene expression analyses, the threshold criteria used to generate clusters accounted for 100% of the unique genes at each temperature.

**Figure 3:**
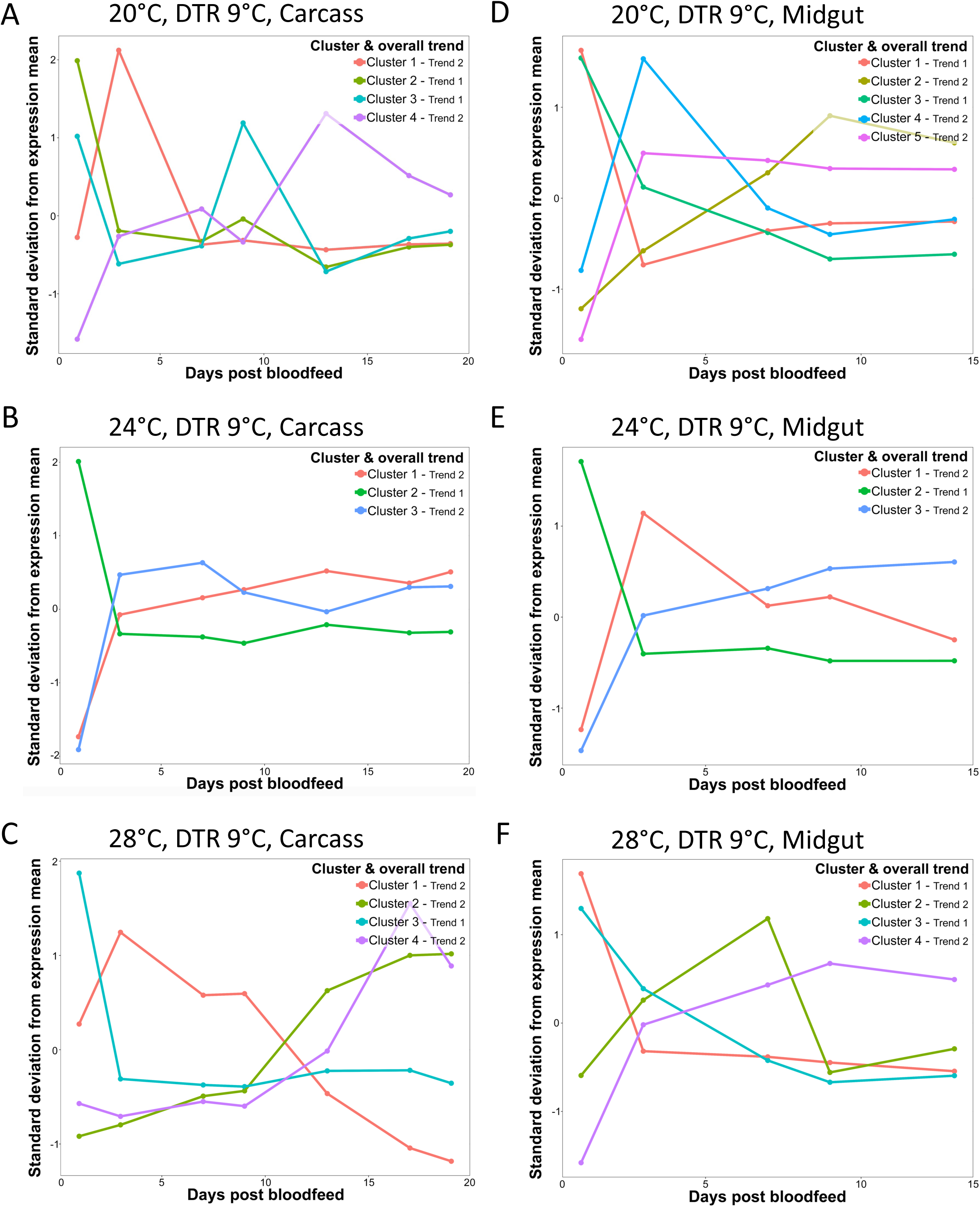
Expression profiles for statistically significant genes, separated by organ and temperature. Each graph represents a different temperature and organ (as labelled), and each line on the graph represents a unique gene expression profile, as determined by Mfuzz and K-means clustering. The colored lines in each panel shows temporal gene expression profiles for each cluster in the carcasses and midguts at 20 (A, D), 24 (B, E) and 28 (C, F) DTR 9°C respectively; x-axes show the sampled time-point in days post-bloodmeal, y-axis shows the standard deviation from the mean expression for that specific gene across the time series (refer to Tables 1 and 2 for a list of the number of genes within a specific cluster). Note the presence of two distinct trends (see key on top-right for each graph): Trend one is marked by high expression of genes one day post-blood meal, followed by subsequent down-regulation, and continued suppression throughout the remaining time-points. Trend two is marked by initial low expression of genes one day post-blood meal, followed by enrichment of genes at subsequent time-points, though the speed and degree of this enrichment is not necessarily consistent.

**Table 1:**
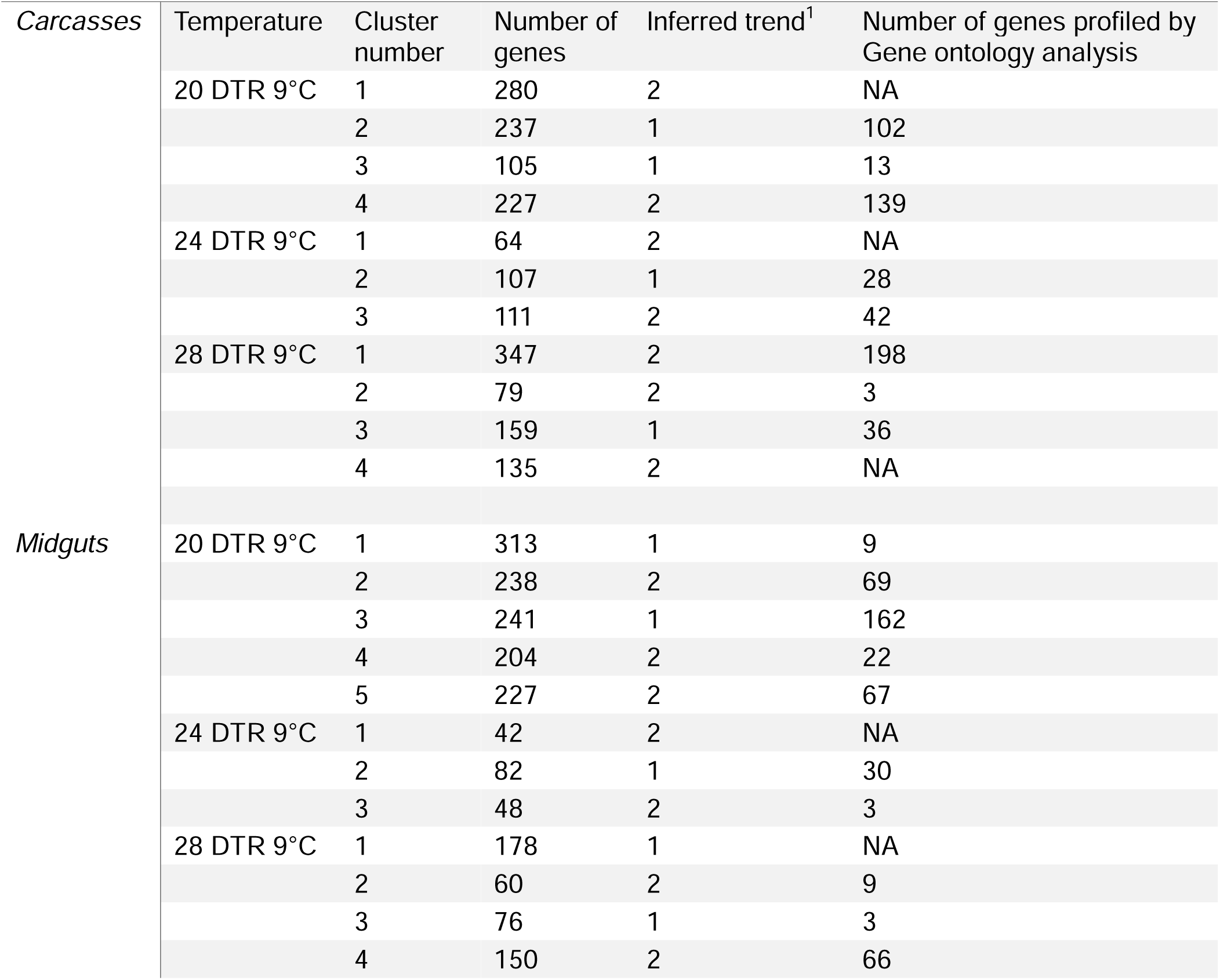
Overview of uniquely expressed gene clusters at each temperature and site of sampling.

**Table 2:** Overview of gene clusters with shared gene expression profiles at each temperature and site of sampling.

| *Carcasses* | Temperature | Cluster number | Number of genes | Inferred trend<sup>1</sup> | Number of genes profiled by Gene ontology analysis |
| --- | --- | --- | --- | --- | --- |
|  | 20 DTR 9°C | 1 | 62 | 2 | 24 |
|  |  | 2 | 70 | 2 | 25 |
|  |  | 3 | 317 | 1 | 192 |
|  | 24 DTR 9°C | 1 | 44 | 2 | 8 |
|  |  | 2 | 24 | 2 | 6 |
|  |  | 3 | 381 | 1 | 219 |
|  | 28 DTR 9°C | 1 | 62 | 2 | 25 |
|  |  | 2 | 92 | 1 | 20 |
|  |  | 3 | 295 | 1 | 162 |
| *Midguts* | 20 DTR 9°C | 1 | 475 | 1 | 227 |
|  |  | 2 | 151 | 2 | 19 |
|  |  | 3 | 165 | 2 | 62 |
|  |  | 4 | 111 | 1 | 52 |
|  | 24 DTR 9°C | 1 | 448 | 1 | 228 |
|  |  | 2 | 125 | 2 | 20 |
|  |  | 3 | 194 | 2 | 143 |
|  |  | 4 | 137 | 1 | 62 |
|  | 28 DTR 9°C | 1 | 385 | 1 | 194 |
|  |  | 2 | 120 | 2 | 2 |
|  |  | 3 | 197 | 2 | 155 |
|  |  | 4 | 200 | 1 | 95 |

Amongst the differentially expressed genes in the carcasses, the expression of 449 genes were shared across all three temperatures (Figure 2B). Grouping these genes based on temporal expression profiles revealed three clusters at each temperature, with majority of the genes falling into cluster 3 (317, 20 DTR 9°C; 381, 24 DTR 9°C; and 295 genes, 28 DTR 9°C; Figure 4A) (Table 2). In the midguts, we observed a higher number of shared genes (902 genes), with 4 clusters at each temperature ranging in size from 111-475 genes with similarities in temporal expression profiles (Figure 4B, Table 2).

**Figure 4:**
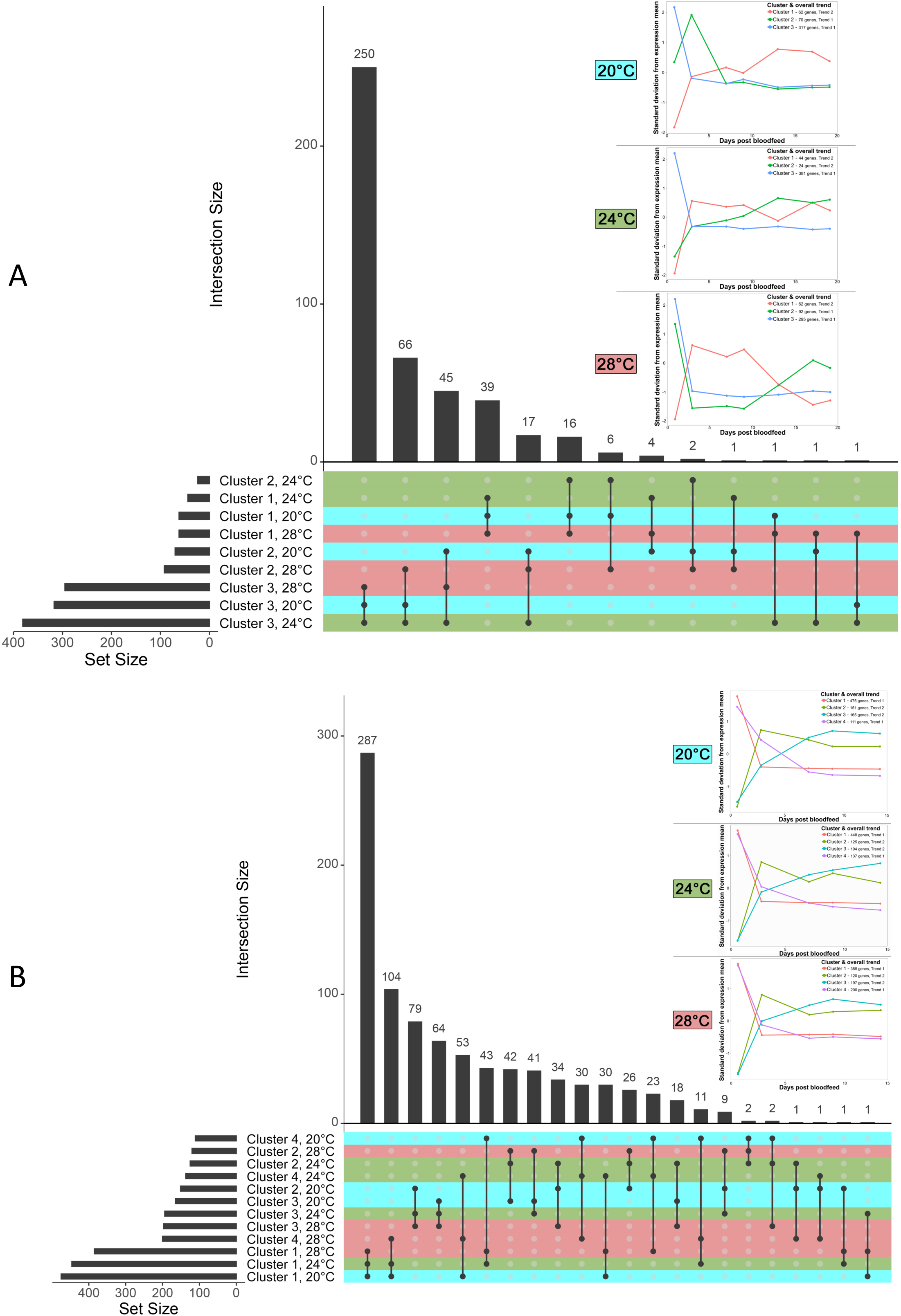
UpsetR plots and expression profiles for genes that showed differential expression at all three temperatures, separated by organ. UpsetR sets are colored based on temperature, which corresponds to the color of the inset label. (3A): Gene expression profiles from carcasses. Note how the largest intersection (250 genes) all correspond to genes from Cluster 3 at the different temperatures (right-most graphs of inset), which are genes that show immediate down-regulation after the first sampled time-point. Subsequent to this, the intersect sizes decreases significantly, with the second-largest set (66 genes) showing genes from Cluster 3 at 20 DTR 9°C and 24 DTR 9°C, with these same genes appearing in Cluster 2 at 28 DTR 9°C, showing enrichment at later time-points. (3B): Gene expression profiles from midguts. Note a more unified pattern of expression profiles, with each temperature having four profiles that show immediate depletion (Cluster 1) or enrichment (Cluster 2) of genes, or a more gradual enrichment (Cluster 3) or depletion (Cluster 4) of genes. The largest intersect (287 genes) in this case comes from Cluster 1 at all three temperatures. The second largest intersect (104) genes show immediate depletion of expression at 20 DTR 9°C and 24 DTR 9°C, but a more gradual rate of depletion at 28 DTR 9°C.

Despite the potential for greater number and diversity of gene expression in the carcasses, variability in gene expression profiles between the various temperatures and time points was less apparent for the carcasses than the midguts, with a larger number of statistically significant genes being observed in midgut tissues. This could be due to a few reasons. The sequencing depth chosen for our study meant only highly expressed genes were retained, with the remaining filtered out during sequencing or by the filtering criteria post-sequencing. Additionally, the presence of other tissues in the carcasses could have resulted in a lower signal to noise ratio, with fewer differentially expressed genes than in mosquito midguts.

### Transcriptomic responses to the blood meal reveals two complementary trends across all gene clusters at both sampling sites and all temperature regimens

A visual comparison of the expression profiles suggested two general trends in expression across the unique and shared gene clusters in the carcasses and midguts at all three temperatures (Figure 3, top-right key and listed in Table 1 in column titled “Inferred trend”). The first trend (Trend 1) was characterized by gene clusters that were enriched a day after the blood meal before returning to expression levels below the averages for the rest of the sampling period. The second trend (Trend 2) by contrast represents gene clusters that were depleted 1 day after the blood meal, but subsequently showed variable expression patterns over the remaining time points, often increasing in expression. Functional profiling of the gene clusters was performed with Gene ontology (GO) enrichment analysis. GO profiling was not possible for all the clusters, especially for some of the uniquely expressed clusters (Table 1) (refer to Supplementary table 1 for detailed output of GO analyses, and Supplementary table 2 for list of protein coding genes corresponding to the GO terms in each cluster). However, the two trends suggest these are genes involved in blood meal digestion, with genes initially induced by the blood meal actively being depleted at the later time points, while genes initially repressed by the blood meal were differentially expressed over the later time points.

#### Functional profiling of Trend 1 genes in carcasses

Amongst the unique gene clusters that were members of Trend 1 (enriched a day after the blood meal, followed by depletion at all other time points) includes cluster 2 at 20 DTR 9°C (237 genes) (Figure 3A, light green line). Analysis of GO enrichment for this profile suggested coordinated regulation of the expression of genes involved in metabolic and catabolic processes of small molecules (primarily organic acids including amino acids, oxoacids, carboxylic acids, and other organic acids). In addition, genes involved in energy generation via respiration (localised primarily in the mitochondria and surrounding membranes) and oxidoreductase functions (e.g., glutathione peroxidase, Supplementary figure 1, Supplementary table 1) were implicated. One exception to the patterns of expression exhibited in Trend 1 at the low temperature treatment was the smaller cluster 3, which showed unusual enrichment at day 9 post-blood meal. GO enrichment for this profile suggested gene products with functions in regulating activity of peptidases involved in inhibiting the activity of serine proteases and metalloproteinases (Supplementary figure 1, Supplementary table 1), which are known to be associated with blood meal digestion, immunity, as well as oogenesis (Hixson et al., 2022; Santiago et al., 2017). In one notable observation, several of the GO terms identified at 20 DTR 9°C have shown enriched gene and protein levels 24h after a blood meal in other studies at higher average constant temperatures of 27-28°C (DTR = 0°C) (Hixson et al., 2022; M. Kumar et al., 2017; Maccallum et al., 2011; Majoline Tchioffo et al., 2021; Short, Mongodin, MacLeod, Talyuli, & Dimopoulos, 2017) but see (Ferreira et al., 2020; Wimalasiri-Yapa et al., 2021). Compared to the two clusters at 20 DTR 9°C (342 genes), only cluster 2 (107 genes) at 24 DTR 9°C (Figure 3B) and cluster 3 (159 genes) at 28 DTR 9°C (Figure 3C) continued to follow Trend 1. GO enrichment of cluster 2 at 24 DTR 9°C suggested genes involved in processes generally associated with localization and transport across membranes for instance, such as monoatomic ions (Supplementary figure 1, Supplementary table 1). Cluster 3 at 28 DTR 9°C also included the iron transport protein transferrin (Supplementary table 2) that was shown to be critical for oogenesis in *An. culicifacies*, with another study demonstrating increased expression in *Aedes aegypti* at 30°C compared to 20°C (Muturi, Blackshear Jr, & Montgomery, 2012; Rani et al., 2022). Taken together, these results indicate that at the cooler temperature of 20 DTR 9°C, some gene products may show delayed/prolonged expression patterns, which could explain the longer gonotrophic cycles in these mosquitoes at cooler temperatures (Miazgowicz et al., 2020). However, they also suggest that diurnal fluctuations may complicate comparisons of gene expression profiles based on average temperatures alone (Breitenbach et al., 2022; Salachan & Sorensen, 2022; Sorensen et al., 2016).

When looking at genes shared across all temperatures, several gene expression profiles appear consistently present (Figure 4A, Table 1). For example, genes in cluster 3 for all three temperatures showed consistent enrichment 1-day post-blood meal, followed by depletion for the remainder of the sampling period. This is despite an apparent delay in blood meal digestion at 20 DTR 9°C. GO annotations suggested that this cluster generally comprised highly enriched genes with products involved in localization and transport (proteins), post-translational modifications (e.g., glycosylation, ufmylation), and processes associated with the endoplasmic reticulum network (Supplementary figure 2). Amongst the GO terms that were highly represented at all three temperatures were biological processes pertaining to ‘signal peptide processing’ and ‘protein ufmylation’, localized to the ‘signal peptidase complex’ and with ‘protein disulfide isomerase activity’ (and the related GO term of ‘intramolecular oxidoreductase activity transposing S-S bonds’) (Supplementary figure 2A, 2B, 2D and Supplementary table 1). Most of the gene products associated with these terms are conserved across phyla and involved in protein folding, degradation, and trafficking to and from the endoplasmic reticulum (Gerakis, Quintero, Li, & Hetz, 2019; Guan et al., 2018). One exception to this is protein disulfide isomerase (Supplementary table 2), which expresses dual oxidoreductase-isomerase functions and serves as an antioxidant in an *Aedes albopictus* cell line infected with DENV (T.-H. Chen et al., 2011), but has also been shown to be essential for blood feeding, survival, and oogenesis in ticks (M. Liao et al., 2008).

#### Functional profiling of Trend 2 genes in carcasses

Genes in the remaining clusters can be grouped into members of Trend 2 (depleted day 1 post-
blood meal, with expression increasing thereafter) and generally displayed more variability in the expression patterns as compared to Trend 1. Amongst the unique gene clusters in the carcasses, seven clusters appeared to follow this trend (Figure 3A-C, see individual keys). For the clusters that GO profiling could be used (Supplementary figure 3), most of the gene products across all three temperatures were generally involved in aspects of nucleic acid (mainly DNA) metabolism, as well as processes localized to the nucleus and capable of binding nucleic acid such as transcription factors (Supplementary table 1). For instance, following depletion at day 1 post-bloodmeal, expression increased over the 19 days for multiple transcription factors, including Nuclear transcription factor Y (20 DTR 9°C, cluster 4), BZIP1 (24 DTR 9°C, cluster 3), and Enhancer of yellow 2 (28 DTR 9°C, cluster 4) (Supplementary table 2). While little is known about the mechanisms of thermoregulation in animals, transcription factors in bacteria and plants play central roles in adjusting physiological responses to temperature changes (Sengupta & Garrity, 2013). While Nuclear transcription factor Y is critical for conferring desiccation tolerance in plants, a recent study suggested its involvement in regulating anhydrobiosis responses in an embryonic cell line from the larvae of the African midge, *Polypedilum vanderplanki* (Yamada et al., 2020). Although the precise function of the BZIP1 transcription factor appears to be unknown, in *Ae. aegypti*, its expression was enriched in the ovaries specifically after a blood meal (Kojin, Biedler, Tu, & Adelman, 2020). Taken together, this result suggests that genes in the clusters that followed Trend 2, were similar in their functional profiles and displayed temperature-dependent shifts in their dynamics of expression. For instance, at the coolest temperature (20 DTR 9°C), expression of the 227 genes in cluster 4 increased until day 13 post-blood meal before declining again (Figure 3A, purple line). At the intermediate temperature (24 DTR 9°C), despite not as pronounced as the cooler temperature, the 111 genes in cluster 3 (Figure 3B, blue line) increased in expression until 7 days post-blood meal before beginning to decline. Finally, at the warmest temperature (28 DTR 9°C), the 347 genes in cluster 1, increased in expression until 3 days post-blood meal before beginning to decline to levels clearly lower than mean expression levels at that temperature by days 17 and 19 post-blood meal (Figure 3C, red line).

Amongst the shared gene clusters in the carcasses, cluster 1 at all three temperatures and cluster 2 at 20 DTR 9°C and 24 DTR 9°C were depleted at day 1 post-blood meal, albeit with differences between the clusters at later time points (Figure 4A). Despite the small size of the clusters (24-70), in general, GO profiling suggests similar functional responses to the unique clusters, with gene products involved in nucleic acid metabolism in the nucleus and the binding of nucleic acids (histones, CDT1 DNA replication factor and DNA helicase) (Supplementary figure 4).

#### Functional profiling of Trend 1 genes in midguts

Amongst the clusters of genes that were uniquely expressed in the midguts at each temperature, genes in clusters 1 and 3 at 20 DTR 9°C and 28 DTR 9°C and cluster 2 at 24 DTR 9°C were characteristic of Trend 1 (Figure 3D-F, see individual keys). Cluster 1 at 20 DTR 9°C primarily comprised genes for transmembrane transporters that utilized ATP for the active transfer of solutes and ions (e.g., calcium-transporting ATPase, Supplementary figure 5A and Supplementary table 1). However, genes in cluster 3 at the same temperature were involved in diverse biological processes such as cellular homeostasis (cell redox homeostasis), carbohydrate metabolism to energy generation (ATP), nucleoside and nucleotide metabolism (including purine synthesis), and other molecular functions to do with cellular respiration and oxidoreductase activity (Supplementary figure 5B). The GO term corresponding to cell redox homeostasis comprised genes coding for glutaredoxin, peroxiredoxin, superoxide dismutase, and protein disulfide isomerase (Supplementary table 2), which are antioxidant proteins that protect against the oxidative environment created by blood meal digestion and *Plasmodium falciparum* infections (Champion & Xu, 2017; Surachetpong, Pakpour, Cheung, & Luckhart, 2010). While GO profiling of genes in cluster 2 at 24 DTR 9°C did not return any robust matches apart from genes with products that bound cyclic compounds such as nucleic acids (Supplementary figure 1C), it was unable to identify any GO terms corresponding to the 178 genes in cluster 1 at 28 DTR 9°C. Analysis of the genes in cluster 3 at the 28 DTR 9°C identified two genes for innexins (Supplementary figure 5D, Supplementary table 2), a family of channel proteins (gap junction and non-junctional) exclusively expressed by invertebrates and essential for intercellular transfer of solutes and ions (Güiza, Barría, Sáez, & Vega, 2018). Genes in the midguts associated with blood meal digestion also show delayed expression at the coolest temperature of 20 DTR 9°C like Trend 1 genes expressed in carcasses.

Amongst the four gene clusters that were shared across the three temperatures, clusters 1 and 4 followed Trend 1, but with gradual instead of immediate depletion of cluster 4 (Figure 4B). For all three temperatures, cluster 1 comprised the largest number of genes with GO profiles reflecting blood meal digestion like catabolism (i.e., breakdown) of proteins, peptides, and functional groups associated with amino acids as part of the proteasome, as well as peptidase complexes for a wide variety of peptidolytic enzymes (Supplementary figure 6A, 6C, 6D and Supplementary table 1). Genes shared across temperatures within cluster 4 were an extension of the previous cluster and included genes with oxidoreductase activities (Supplementary figure 1B, 1E, 1F). Cluster 4 at 28 DTR 9°C was larger than the other Cluster 4 (200 genes) (Supplementary figure 1F) with GO analysis predicting the additional gene products to have peptidase functions and iron binding activities likely due to the breakdown of hemoglobin during blood meal digestion. This is in addition to ferritin and heme oxygenase activity (Supplementary table 2), which was previously shown to be required for oogenesis in *An. gambiae* (Spencer et al., 2018).

#### Functional profiling of Trend 2 genes in midguts

Across all three temperatures in the midguts, a total of 7 gene clusters can be considered members of Trend 2, with initial depletion followed by varying cluster-dependent trends in expression at later time-points. For example, depletion of the 238 genes in cluster 2 at 20 DTR 9°C one day after the blood meal was replaced by a sustained increase in expression levels until 13 days post-blood meal (Figure 3D, light green line). Subsequent GO profiling suggested genes in this cluster were primarily associated with RNA synthesis and transcription in the nucleus (Supplementary figure 7A) (Supplementary table 1). In contrast, while the expression of genes in cluster 4 (204 genes) and cluster 5 (227 genes) at 20°C showed a sharp increase at day 3 post-blood meal (Figure 3), only genes in cluster 4 were expressed at below-average levels for the remaining time-points. While GO profiling was unable to identify clear functional associations for cluster 4, it suggested gene products in cluster 5 were associated with loading tRNA for protein translation as well as processing non-coding RNA (ncRNA) (Supplementary figure 7C). The importance of ncRNA in regulating gene expression and facilitating rapid responses to environmental change in mosquitoes is increasingly being recognized (Farley, Eggleston, & Riehle, 2021). At 24 DTR 9°C, clusters 1 and 3 were also members of Trend 2 (Figure 3B). While GO profiling was unable to determine functional associations between the genes in cluster 1, only 3 genes in cluster 3 could be identified with roles in detection and response to external stimuli, potentially for finding nutrition, vertebrate hosts, and oviposition sites (Peach & Blake, 2023). At 28 DTR 9°C, clusters 2 and 4 were identified as members of Trend 2. GO profiling of genes in cluster 2 identified roles in regulating peptidase activity, while genes in cluster 4 were associated with initialization and regulation of protein translation (e.g., Eukaryotic translation initiation factor 3), and localization to the ribosomes (e.g., 40S ribosomal proteins S7 and SA) (Supplementary table 2). The identification of transcripts for 40S ribosomal protein S7 increasing in expression over time, specifically at 28 DTR 9°C, indicates that caution should be used against relying on this gene as a reference/housekeeping gene (Calkins & Piermarini, 2017; Kojin et al., 2020; Liu et al., 2022; Roy et al., 2022).

Amongst clusters shared across all three temperatures, genes in clusters 2 and 3 were identified as members of Trend 2 (Figure 4B). Genes in cluster 2 were primarily associated with protein translation and folding and showed rapid increases by day 3 post-blood meal before returning to levels close to the average for the duration of the study. While the products of genes in cluster 3 were also associated with translation and protein folding, their expression gradually increased over the 13 days (Supplementary figure 8) (Supplementary table 1).

## CONCLUSIONS

Our study has several important implications. First, we found most of the variation in gene expression was dependent on the tissue sampled (36% of the overall variance), demonstrating clear spatial regulation of gene expression in response to the blood meal (25% of the overall variation). We identified differentially expressed genes that were shared across all three temperatures, suggesting that the expression of these genes was critical to regulating various processes of mosquito life history. Second, of the differentially expressed genes that were unique to a given temperature, our results suggest that mosquitoes may tolerate cool temperature compared to warm temperature fluctuations by expressing a larger and more diverse repertoire of gene products. Third, we noticed that clusters of genes tended to display similar gene expression patterns over time, and these patterns followed two general trends representing a coordinated response to blood meal digestion, managing oxidative stress, and channeling resources from the blood meal towards reproduction. These temporal dynamics were also temperature sensitive, reinforcing the many studies done to date that demonstrate the effects of temperature variation on various aspects of the mosquito life cycle, such as egg to adult development rates, lifespan, fecundity, biting rate as well as vectorial capacity (reviewed by (Miazgowicz et al., 2020; Mordecai et al., 2019)). However, our results also suggest that incorporating more realistic diurnal temperature fluctuations may influence gene expression profiles in distinct ways than constant temperature variation. Finally, as one of the more comprehensive examinations of temporal variation in mosquito gene expression, our study also cautions against the use of 40S ribosomal protein S7 as a reference / housekeeping gene for relative quantification of gene expression, as its expression violates the assumption that its expression is insensitive to experimental perturbations.

## ACKNOWLEDGEMENTS

Funding for this work was provided by the University of Georgia and the NIH NIAID grants (5R01AI110793-04, R01 AI163444-01). The funding bodies did not have any say in the design of the study and collection, analysis, interpretation of data or in writing the manuscript. AKP was also supported by the Georgia Research Alliance, the SporoCore and 5P30AI168386-02 (sub-award A866298). GLH was supported by the BBSRC (BB/V011278/1, BB/X018024/1, and BB/W018446/1), the UKRI (20197), a Royal Society Wolfson Fellowship (RSWF\R1\180013), the NIHR (NIHR2000907), and the Bill and Melinda Gates Foundation (INV-048598).

## DATA ACCESSIBILITY

Raw RNA sequencing datasets have been deposited in NCBI, under BioProject ID Number PRJNA1116119. All supplementary data has been included as additional data tables with submission.

## AUTHOR CONTRIBUTIONS

Designed research: AKP, CCM.

Performed Research: AKP, RS, JCS.

Contributed new reagents or analytical tools: MBT, GLH.

Analyzed data: SQ, GLH.

Wrote the paper: AKP, SQ, CCM.

Funding acquisition: CCM, MBT, GLH.

## FIGURE AND TABLE CAPTIONS

**Supplementary figure 1:**
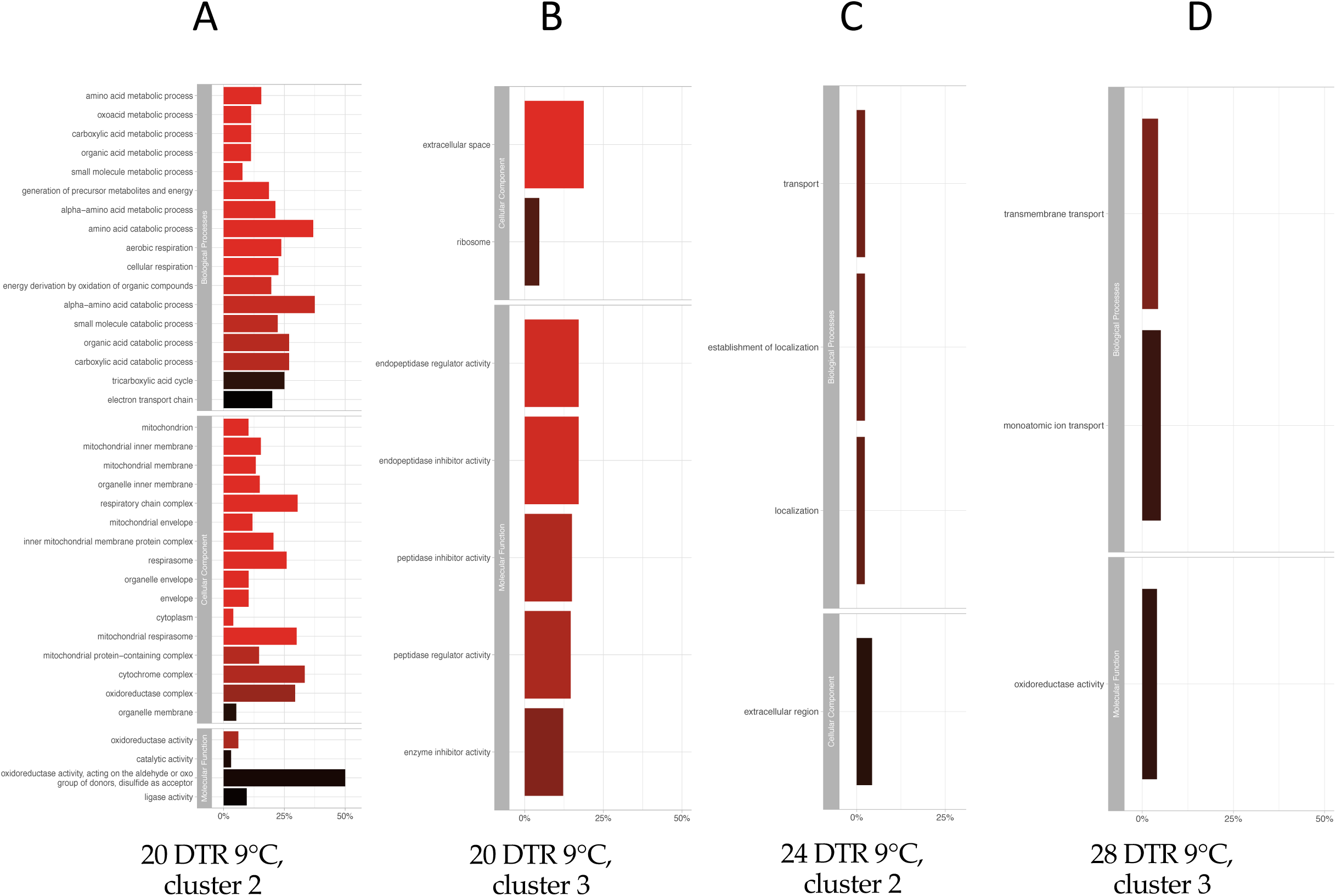
GO profiles of Unique gene clusters in carcasses following Trend 1: enriched clusters at 1 dpi and then reduced thereafter. The GO enrichment graphs show the top five GO categories within the three major branches of the GO domains (‘Biological Process’, ‘Cellular Component’ and ‘Molecular Function’). The y-axis of these graphs indicates the GO term, the x-axis represents the percentage of gene products associated with the GO term, and the color of the bar represents the statistical significance of enrichment (with brighter shading indicating greater significance).

**Supplementary figure 2:**
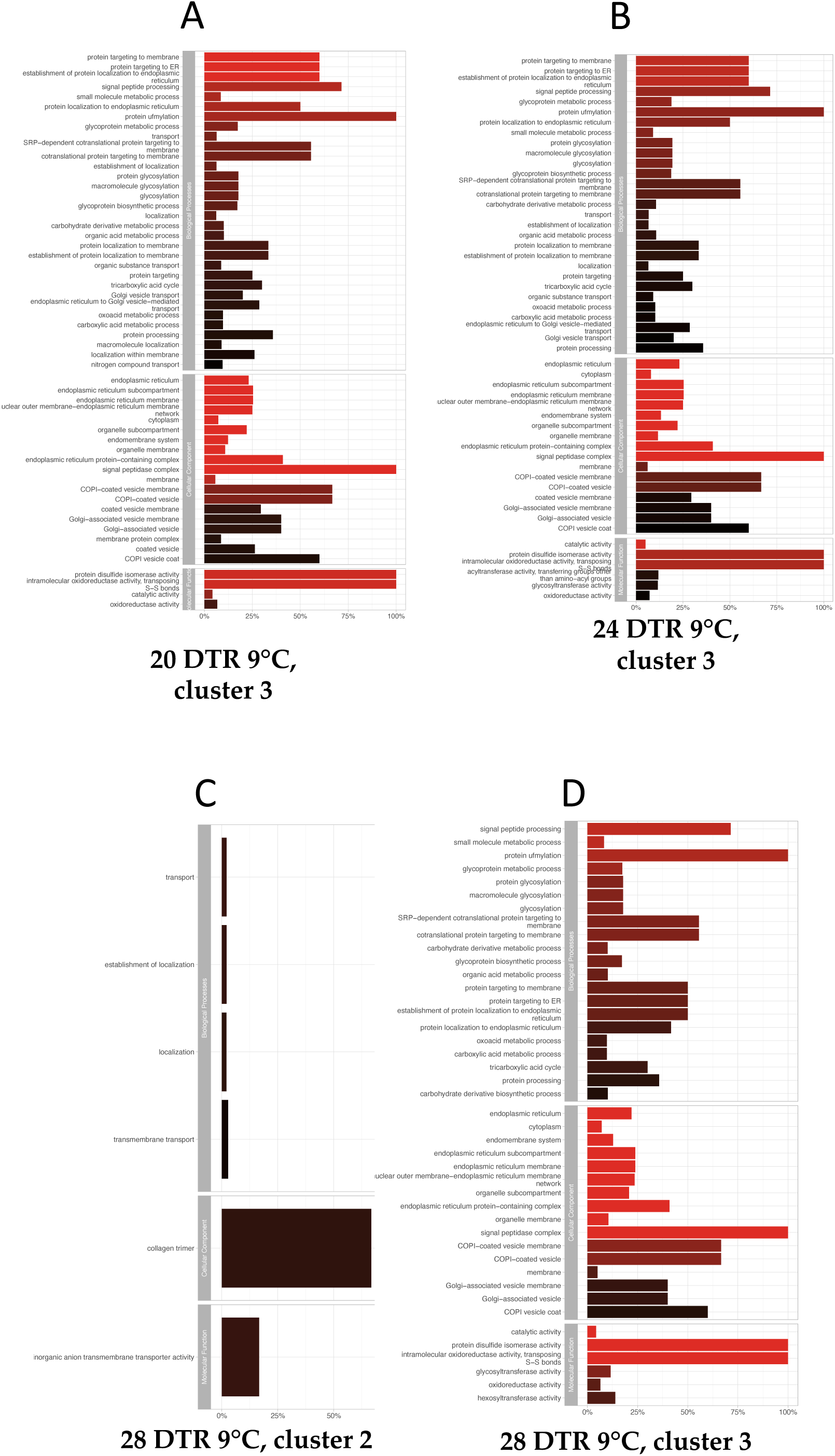
GO profiles of shared gene clusters in carcasses following Trend 1: enriched clusters at 1dpi and then reduced thereafter.

**Supplementary figure 3:**
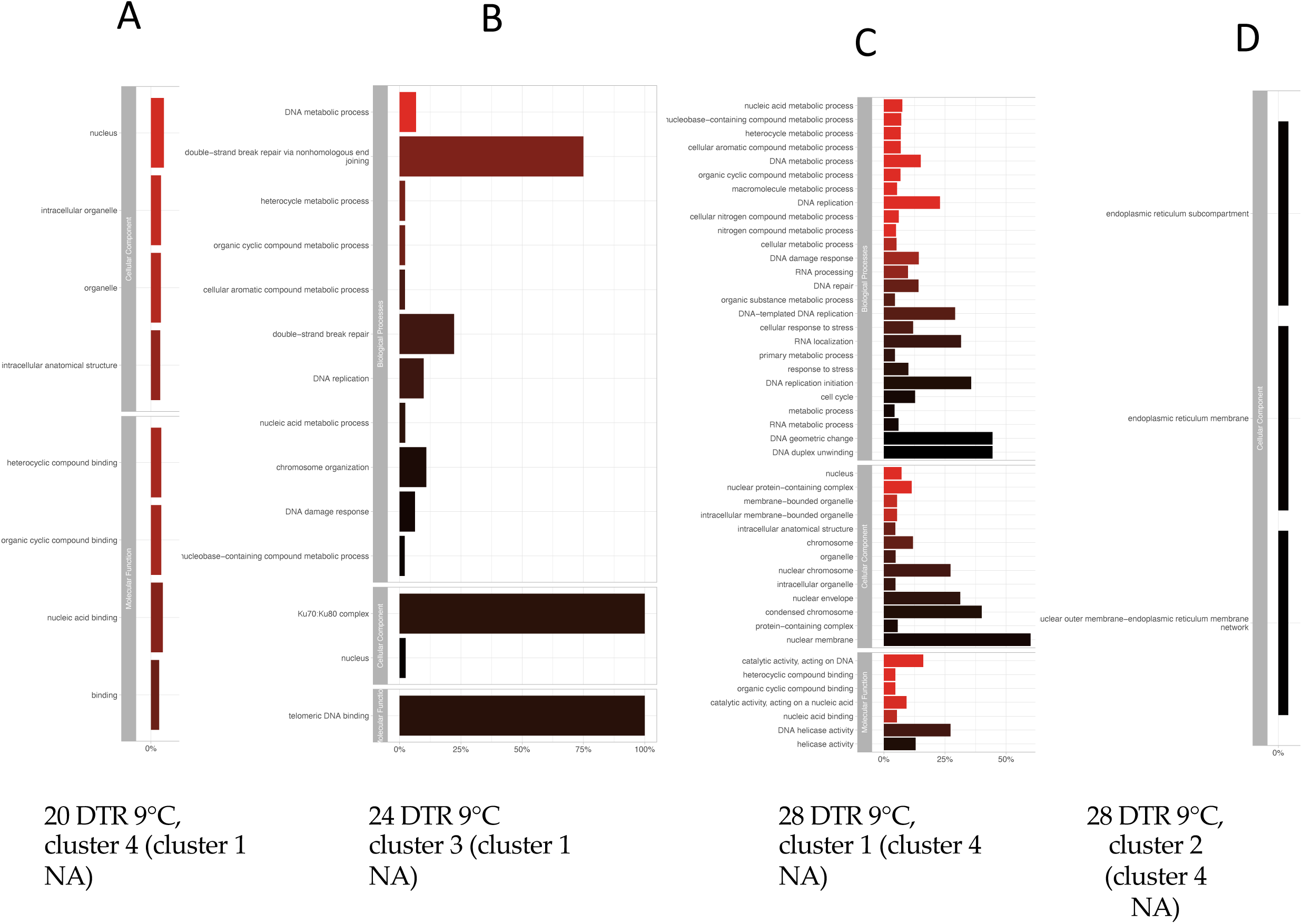
GO profiles of unique gene clusters in carcasses following Trend 2: depleted clusters at 1dpi, variable thereafter.

**Supplementary figure 4:**
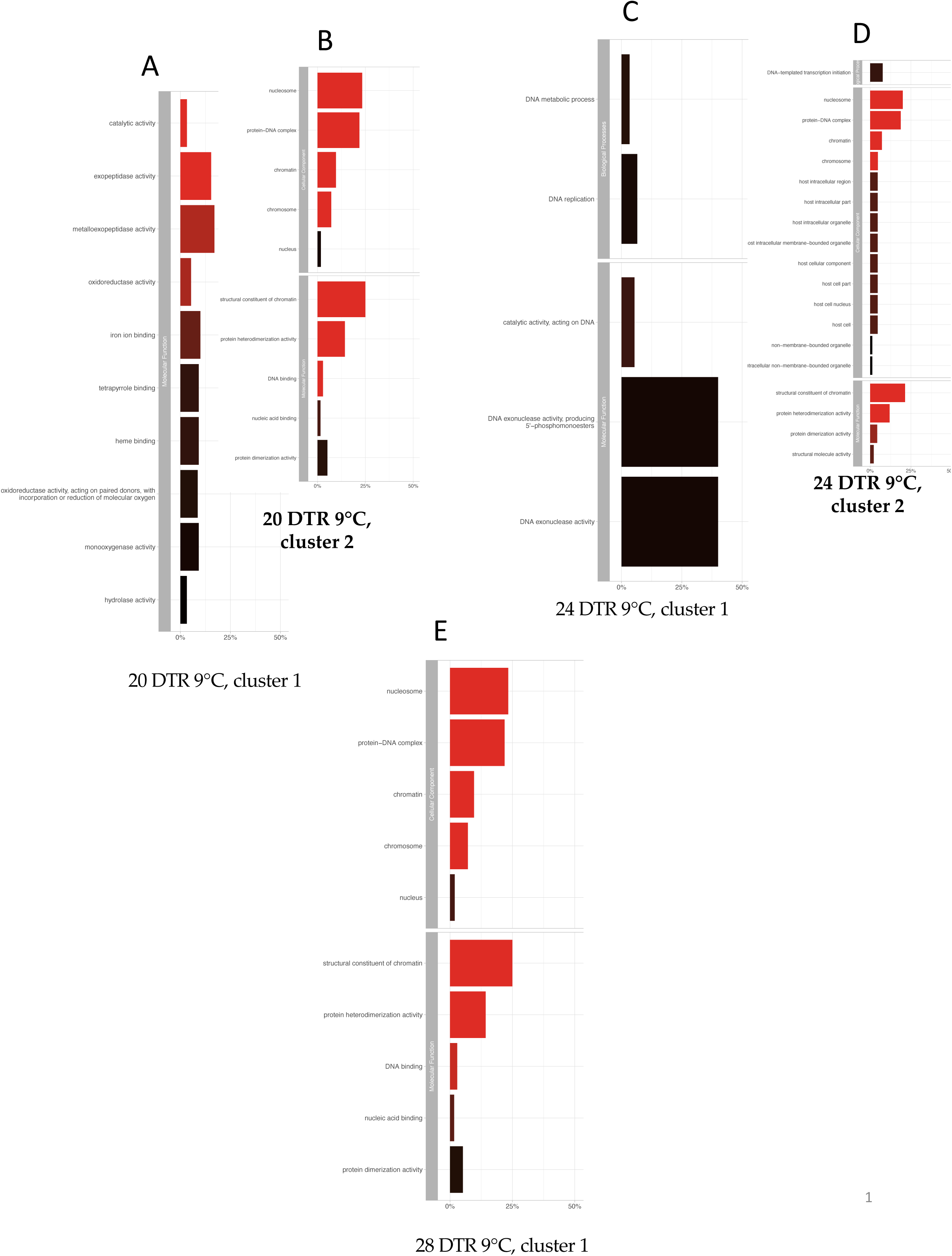
GO profiles of shared gene clusters in carcasses following Trend 2: depleted clusters at 1dpi, variable thereafter.

**Supplementary figure 5:**
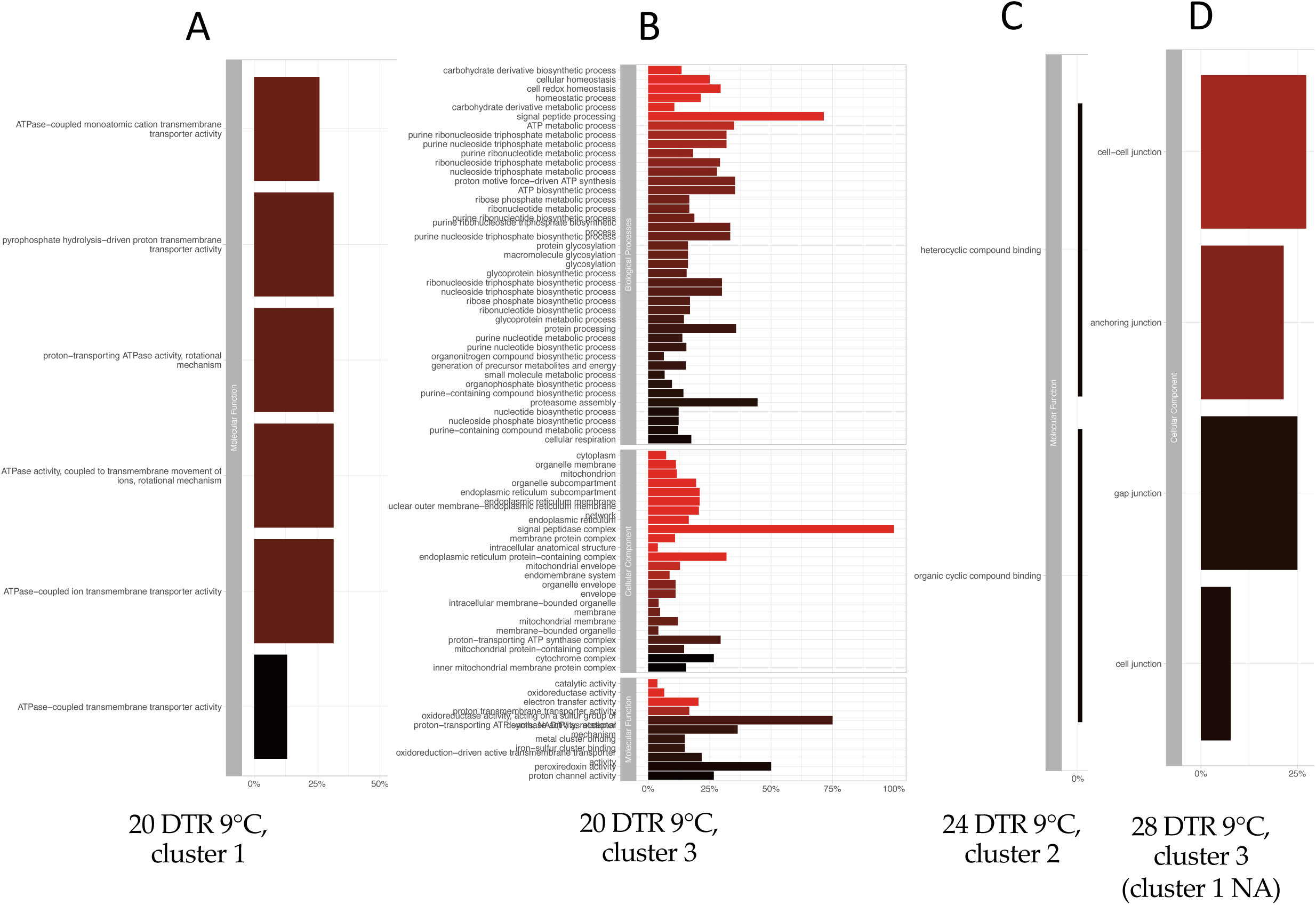
GO profiles of unique gene clusters in midguts following Trend 1: enriched clusters at 1dpi and then reduced thereafter.

**Supplementary figure 6:**
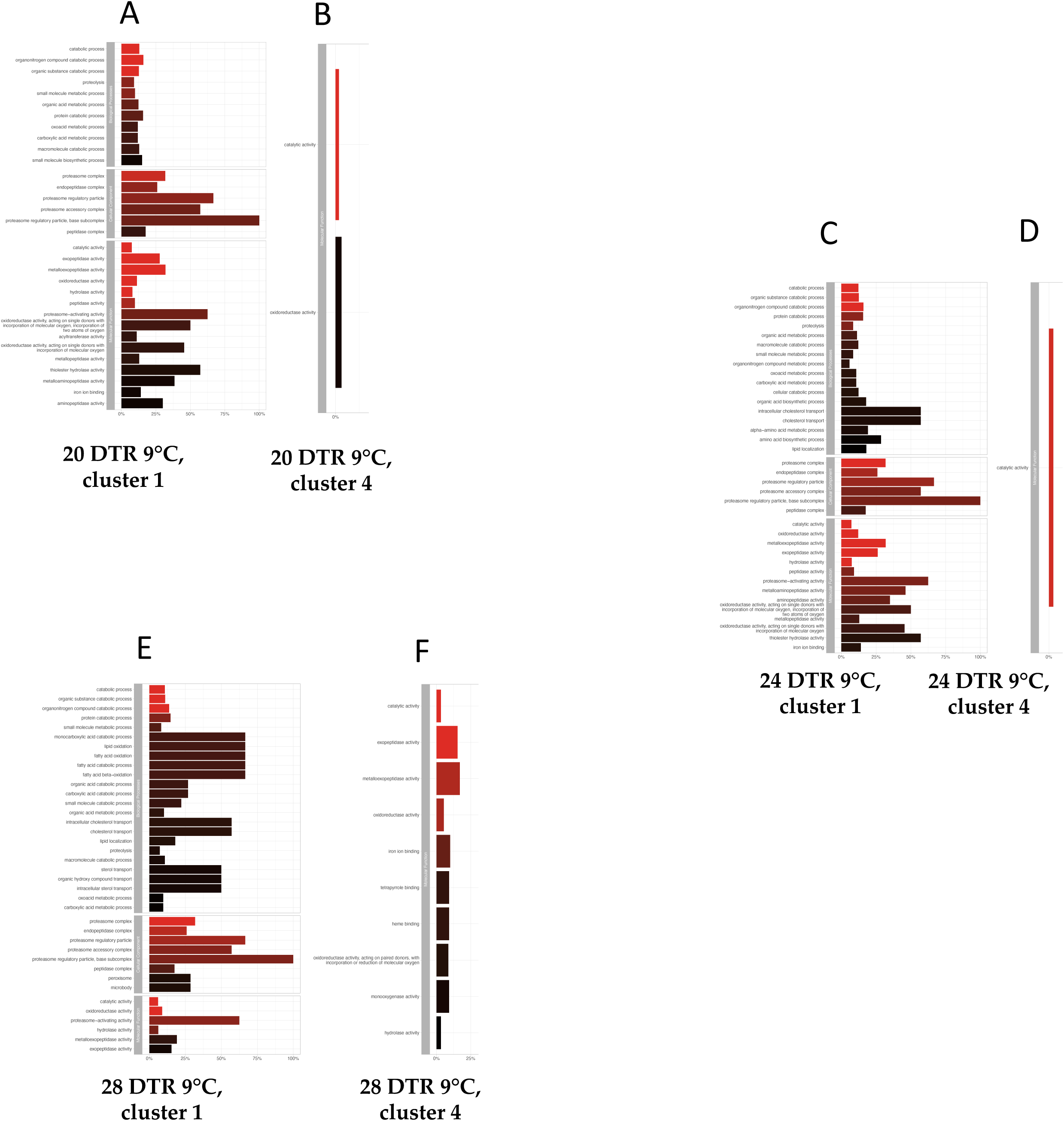
GO profiles of shared gene clusters in midguts following Trend 1: enriched clusters at 1dpi and then reduced thereafter.

**Supplementary figure 7:**
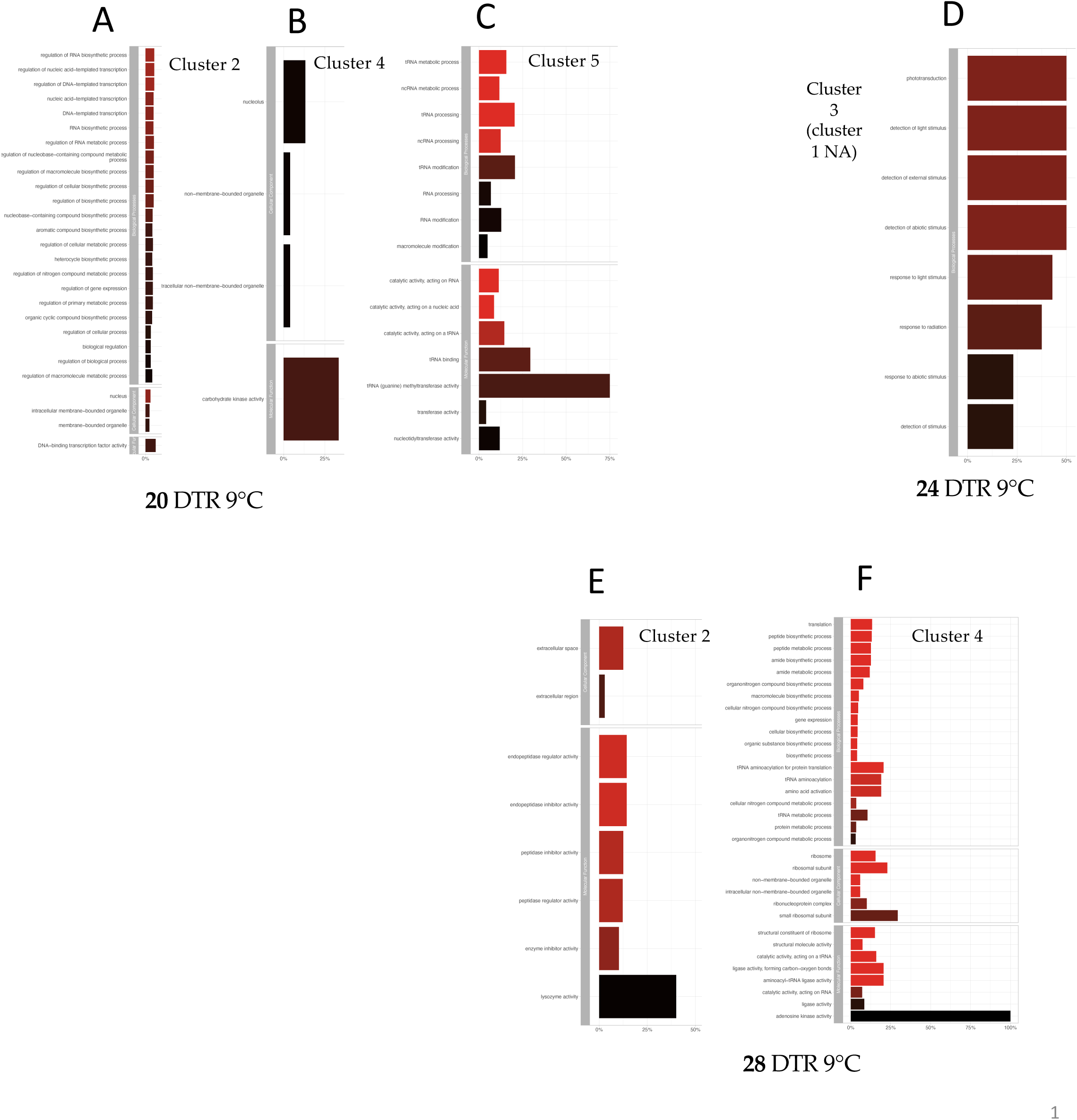
GO profiles of unique gene clusters in midguts following Trend 2: depleted clusters at 1dpi, variable thereafter.

**Supplementary figure 8:**
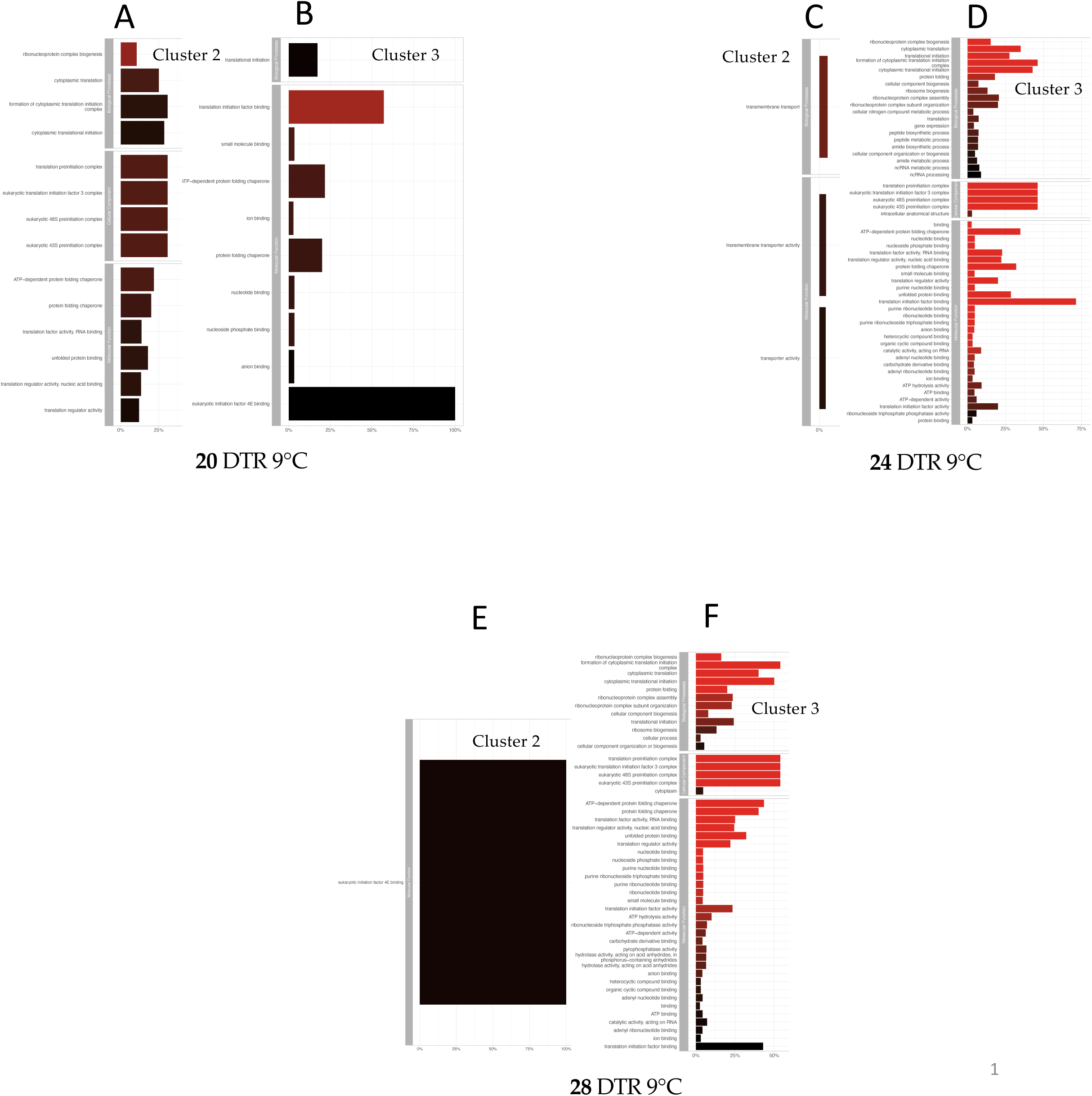
GO profiles of shared gene clusters in midguts following Trend 2: depleted clusters at 1dpi, variable thereafter.

**Supplementary table 1:** Results of GO profiling grouped and sortable according to cluster type (unique or shared. column titled ‘Cluster.type’), site of sampling (carcasses and midguts, column titled ‘Tissue’), temperature, (20, 24 and 28 DTR 9°C, column ‘Temperature’) and finally the cluster number (column ‘Cluster.no’).

**Supplementary table 2:** List of gene products corresponding to the GO profiles in Supplementary table 1. Results are also grouped and sortable in the same format as Supplementary table 1.

## REFERENCES

Andrews, S. (2010). FastQC: A Quality Control Tool for High Throughput Sequence Data [Online]. http://www.bioinformatics.babraham.ac.uk/projects/fastqc/. Retrieved from http://www.bioinformatics.babraham.ac.uk/projects/fastqc/

Breitenbach, A. T., Bowden, R. M., & Paitz, R. T. (2022). Effects of Constant and Fluctuating Temperatures on Gene Expression During Gonadal Development. Integrative and Comparative Biology, 62(1), 21–29. doi:10.1093/icb/icac011

Brown, J. J., Pascual, M., Wimberly, M. C., Johnson, L. R., & Murdock, C. C. (2023). Humidity - The overlooked variable in the thermal biology of mosquito-borne disease. Ecol Lett, 26(7), 1029–1049. doi:10.1111/ele.14228

Calkins, T. L., & Piermarini, P. M. (2017). A Blood Meal Enhances Innexin mRNA Expression in the Midgut, Malpighian Tubules, and Ovaries of the Yellow Fever Mosquito Aedes aegypti. Insects, 8(4). doi:10.3390/insects8040122

Cassone, B. J., Molloy, M. J., Cheng, C., Tan, J. C., Hahn, M. W., & Besansky, N. J. (2011). Divergent transcriptional response to thermal stress by Anopheles gambiae larvae carrying alternative arrangements of inversion 2La. Molecular Ecology, 20(12), 2567–2580. 10.1111/j.1365-294X.2011.05114.x

Champion, C. J., & Xu, J. (2017). The impact of metagenomic interplay on the mosquito redox homeostasis. Free Radical Biology and Medicine, 105, 79–85. 10.1016/j.freeradbiomed.2016.11.031

Chen, J. L., & Lewis, O. T. (2024). Limits to species distributions on tropical mountains shift from high temperature to competition as elevation increases. ECOLOGICAL MONOGRAPHS, 94(1). doi:10.1002/ecm.1597

Chen, T.-H., Tang, P., Yang, C.-F., Kao, L.-H., Lo, Y.-P., Chuang, C.-K., … Chen, W.-J. (2011). Antioxidant defense is one of the mechanisms by which mosquito cells survive dengue 2 viral infection. Virology, 410(2), 410–417. 10.1016/j.virol.2010.12.013

Conway, J. R., Lex, A., & Gehlenborg, N. (2017). UpSetR: an R package for the visualization of intersecting sets and their properties. Bioinformatics, 33(18), 2938–2940. doi:10.1093/bioinformatics/btx364

Deutsch, C. A., Tewksbury, J. J., Huey, R. B., Sheldon, K. S., Ghalambor, C. K., Haak, D. C., & Martin, P. R. (2008). Impacts of climate warming on terrestrial ectotherms across latitude. Proceedings of the National Academy of Sciences, 105(18), 6668–6672. doi:10.1073/pnas.0709472105

Domingos, A., Pinheiro-Silva, R., Couto, J., do Rosario, V., & de la Fuente, J. (2017). The Anopheles gambiae transcriptome - a turning point for malaria control. Insect Mol Biol, 26(2), 140–151. doi:10.1111/imb.12289

Farley, E. J., Eggleston, H., & Riehle, M. M. (2021). Filtering the Junk: Assigning Function to the Mosquito Non-Coding Genome. Insects, 12(2). Retrieved from doi:10.3390/insects12020186

Ferreira, P. G., Tesla, B., Horacio, E. C. A., Nahum, L. A., Brindley, M. A., de Oliveira Mendes, T. A., & Murdock, C. C. (2020). Temperature Dramatically Shapes Mosquito Gene Expression With Consequences for Mosquito-Zika Virus Interactions. Front Microbiol, 11, 901. doi:10.3389/fmicb.2020.00901

Gerakis, Y., Quintero, M., Li, H., & Hetz, C. (2019). The UFMylation System in Proteostasis and Beyond. Trends in Cell Biology, 29(12), 974–986. doi:10.1016/j.tcb.2019.09.005

González-Tokman, D., Córdoba-Aguilar, A., Dáttilo, W., Lira-Noriega, A., Sánchez-Guillén, R. A., & Villalobos, F. (2020). Insect responses to heat: physiological mechanisms, evolution and ecological implications in a warming world. BIOLOGICAL REVIEWS, 95(3), 802–821. doi:10.1111/brv.12588

Guan, J., Zhang, J., Yuan, S., Yang, B., Clark, K. D., Ling, E., & Huang, W. (2018). Analysis of the functions of the signal peptidase complex in the midgut of Tribolium castaneum. Archives of Insect Biochemistry and Physiology, 97(3), e21441. 10.1002/arch.21441

Güiza, J., Barría, I., Sáez, J. C., & Vega, J. L. (2018). Innexins: Expression, Regulation, and Functions. Frontiers in Physiology, 9.

Harvey, J. A., Tougeron, K., Gols, R., Heinen, R., Abarca, M., Abram, P. K., … Chown, S. L. (2023). Scientists’ warning on climate change and insects. ECOLOGICAL MONOGRAPHS, 93(1). doi:10.1002/ecm.1553

Hixson, B., Bing, X.-L., Yang, X., Bonfini, A., Nagy, P., & Buchon, N. (2022). A transcriptomic atlas of Aedes aegypti reveals detailed functional organization of major body parts and gut regional specializations in sugar-fed and blood-fed adult females. Elife, 11, e76132. doi:10.7554/eLife.76132

Johnson, L. R., Ben-Horin, T., Lafferty, K. D., McNally, A., Mordecai, E., Paaijmans, K. P., … Ryan, S. J. (2015). Understanding uncertainty in temperature effects on vector-borne disease: a Bayesian approach. Ecology, 96(1), 203–213. doi:10.1890/13-1964.1

Kim, D., Paggi, J. M., Park, C., Bennett, C., & Salzberg, S. L. (2019). Graph-based genome alignment and genotyping with HISAT2 and HISAT-genotype. Nat Biotechnol, 37(8), 907–915. doi:10.1038/s41587-019-0201-4

Kingsolver, J. G., & Buckley, L. B. (2017). Quantifying thermal extremes and biological variation to predict evolutionary responses to changing climate. PHILOSOPHICAL TRANSACTIONS OF THE ROYAL SOCIETY B-BIOLOGICAL SCIENCES, 372(1723). doi:10.1098/rstb.2016.0147

Kirk, D., O’Connor, M. I., & Mordecai, E. A. (2022). Scaling effects of temperature on parasitism from individuals to populations. J Anim Ecol, 91(10), 2087–2102. doi:10.1111/1365-2656.13786

Kojin, B. B., Biedler, J. K., Tu, Z., & Adelman, Z. N. (2020). Characterization of a female germline and early zygote promoter from the transcription factor bZip1 in the dengue mosquito Aedes aegypti. Parasites & Vectors, 13(1), 353. doi:10.1186/s13071-020-04216-w

Kumar, L., & M, E. F. (2007). Mfuzz: a software package for soft clustering of microarray data. Bioinformation, 2(1), 5–7. doi:10.6026/97320630002005

Kumar, M., Mohanty, A. K., Sreenivasamurthy, S. K., Dey, G., Advani, J., Pinto, S. M., … Prasad, T. S. K.. (2017). Response to Blood Meal in the Fat Body of Anopheles stephensi Using Quantitative Proteomics: Toward New Vector Control Strategies Against Malaria. OMICS, 21(9), 520–530. doi:10.1089/omi.2017.0092

Liao, M., Boldbaatar, D., Gong, H., Huang, P., Umemiya, R., Harnnoi, T., … Fujisaki, K. (2008). Functional analysis of protein disulfide isomerases in blood feeding, viability and oocyte development in Haemaphysalis longicornis ticks. Insect Biochemistry and Molecular Biology, 38(3), 285–295. 10.1016/j.ibmb.2007.11.006

Liao, Y., Smyth, G. K., & Shi, W. (2014). featureCounts: an efficient general purpose program for assigning sequence reads to genomic features. Bioinformatics, 30(7), 923–930. doi:10.1093/bioinformatics/btt656

Liu, Z., Xu, Y., Li, Y., Xu, S., Li, Y., Xiao, L., … Zheng, K. (2022). Transcriptome analysis of Aedes albopictus midguts infected by dengue virus identifies a gene network module highly associated with temperature. Parasites & Vectors, 15(1), 173. doi:10.1186/s13071-022-05282-y

Logan, M. L., & Cox, C. L. (2020). Genetic Constraints, Transcriptome Plasticity, and the Evolutionary Response to Climate Change. Front Genet, 11, 538226. doi:10.3389/fgene.2020.538226

Love, M. I., Huber, W., & Anders, S. (2014). Moderated estimation of fold change and dispersion for RNA-seq data with DESeq2. Genome Biol, 15(12), 550. doi:10.1186/s13059-014-0550-8

Maccallum, R. M., Redmond, S. N., & Christophides, G. K. (2011). An expression map for Anopheles gambiae. BMC Genomics, 12(1471-2164 (Electronic)), 620. doi:10.1186/1471-2164-12-620

Majoline Tchioffo, T., Etienne, K., Nicolas, P., Solomon, E., Caroline, P., Jessy, G.-Y., … Catherine, B. (2021). Differential transcriptomic response of *Anopheles arabiensis* to *Plasmodium vivax* and *Plasmodium falciparum infection*. bioRxiv, 2021.2005.2028.446219. doi:10.1101/2021.05.28.446219

Miazgowicz, K. L., Shocket, M. S., Ryan, S. J., Villena, O. C., Hall, R. J., Owen, J., … Murdock, C. C. (2020). Age influences the thermal suitability of Plasmodium falciparum transmission in the Asian malaria vector Anopheles stephensi. Proc Biol Sci, 287(1931), 20201093. doi:10.1098/rspb.2020.1093

Mordecai, E. A., Caldwell, J. M., Grossman, M. K., Lippi, C. A., Johnson, L. R., Neira, M., … Villena, O. (2019). Thermal biology of mosquito-borne disease. Ecol Lett, 22(10), 1690–1708. doi:10.1111/ele.13335

Murdock, C. C., Blanford, S., Luckhart, S., & Thomas, M. B. (2014). Ambient temperature and dietary supplementation interact to shape mosquito vector competence for malaria. J Insect Physiol, 67, 37–44. doi:10.1016/j.jinsphys.2014.05.020

Murdock, C. C., Sternberg, E. D., & Thomas, M. B. (2016). Malaria transmission potential could be reduced with current and future climate change. Sci Rep, 6, 27771. doi:10.1038/srep27771

Muturi, E. J., Blackshear Jr, M., & Montgomery, A. (2012). Temperature and density-dependent effects of larval environment on Aedes aegypti competence for an alphavirus. Journal of Vector Ecology, 37(1), 154–161. 10.1111/j.1948-7134.2012.00212.x

Oomen, R. A., & Hutchings, J. A. (2022). Genomic reaction norms inform predictions of plastic and adaptive responses to climate change. J Anim Ecol, 91(6), 1073–1087. doi:10.1111/1365-2656.13707

Pathak, A. K., Shiau, J. C., Thomas, M. B., & Murdock, C. C. (2018). Cryogenically preserved RBCs support gametocytogenesis of Plasmodium falciparum in vitro and gametogenesis in mosquitoes. Malar J, 17(1), 457. doi:10.1186/s12936-018-2612-y

Pathak, A. K., Shiau, J. C., Thomas, M. B., & Murdock, C. C. (2019). Field Relevant Variation in Ambient Temperature Modifies Density-Dependent Establishment of Plasmodium falciparum Gametocytes in Mosquitoes. Front Microbiol, 10, 2651. doi:10.3389/fmicb.2019.02651

Peach, D. A. H., & Blake, A. J. (2023). Mosquito (Diptera: Culicidae) Vision and Associated Electrophysiological Techniques. Cold Spring Harb Protoc, 2023(10), pdb top107671. doi:10.1101/pdb.top107671

R Core Team. (2021). R: A Language and Environment for Statistical Computing. Vienna, Austria: R Foundation for Statistical Computing. Retrieved from https://www.R-project.org

Rani, J., De, T. D., Chauhan, C., Kumari, S., Sharma, P., Tevatiya, S., … Dixit, R. (2022). Functional disruption of transferrin expression alters reproductive physiology in Anopheles culicifacies. PLoS One, 17(3), e0264523. doi:10.1371/journal.pone.0264523

Reimand, J., Kull, M., Peterson, H., Hansen, J., & Vilo, J. (2007). g:Profiler--a web-based toolset for functional profiling of gene lists from large-scale experiments. Nucleic Acids Res, 35(Web Server issue), W193–200. doi:10.1093/nar/gkm226

Roy, S., Saha, T. T., Ha, J., Banerjee, R., Aksoy, E., Kulkarni, A., & Raikhel, A. S. (2022). Direct and indirect gene repression by the ecdysone cascade during mosquito reproductive cycle. Proc Natl Acad Sci U S A, 119(11), e2116787119. doi:10.1073/pnas.2116787119

Salachan, P. V., & Sorensen, J. G. (2022). Molecular mechanisms underlying plasticity in a thermally varying environment. Mol Ecol, 31(11), 3174–3191. doi:10.1111/mec.16463

Santiago, P. B., de Araújo, C. N., Motta, F. N., Praça, Y. R., Charneau, S., Bastos, I. M. D., & Santana, J. M. (2017). Proteases of haematophagous arthropod vectors are involved in blood-feeding, yolk formation and immunity - a review. Parasites & Vectors, 10(1), 79. doi:10.1186/s13071-017-2005-z

Sengupta, P., & Garrity, P. (2013). Sensing temperature. Curr Biol, 23(8), R304–307. doi:10.1016/j.cub.2013.03.009

Shapiro, L. L. M., Whitehead, S. A., & Thomas, M. B. (2017). Quantifying the effects of temperature on mosquito and parasite traits that determine the transmission potential of human malaria. PLoS Biol, 15(10), e2003489. doi:10.1371/journal.pbio.2003489

Short, S. M., Mongodin, E. F., MacLeod, H. J., Talyuli, O. A. C., & Dimopoulos, G. (2017). Amino acid metabolic signaling influences Aedes aegypti midgut microbiome variability. PLoS Negl Trop Dis, 11(7), e0005677. doi:10.1371/journal.pntd.0005677

Sinka, M. E., Pironon, S., Massey, N. C., Longbottom, J., Hemingway, J., Moyes, C. L., & Willis, K. J. (2020). A new malaria vector in Africa: Predicting the expansion range of Anopheles stephensi and identifying the urban populations at risk. Proc Natl Acad Sci U S A, 117(40), 24900–24908. doi:10.1073/pnas.2003976117

Sorensen, J. G., Schou, M. F., Kristensen, T. N., & Loeschcke, V. (2016). Thermal fluctuations affect the transcriptome through mechanisms independent of average temperature. Sci Rep, 6, 30975. doi:10.1038/srep30975

Spencer, C. S., Yunta, C., de Lima, G. P. G., Hemmings, K., Lian, L. Y., Lycett, G., & Paine, M. J. I. (2018). Characterisation of Anopheles gambiae heme oxygenase and metalloporphyrin feeding suggests a potential role in reproduction. Insect Biochem Mol Biol, 98(1879-0240 (Electronic)), 25–33. doi:10.1016/j.ibmb.2018.04.010

Sunday, J. M., Bates, A. E., & Dulvy, N. K. (2012). Thermal tolerance and the global redistribution of animals. NATURE CLIMATE CHANGE, 2(9), 686–690. doi:10.1038/NCLIMATE1539

Surachetpong, W., Pakpour, N., Cheung, K. W., & Luckhart, S. (2010). Reactive Oxygen Species-Dependent Cell Signaling Regulates the Mosquito Immune Response to Plasmodium falciparum. Antioxidants & Redox Signaling, 14(6), 943–955. doi:10.1089/ars.2010.3401

Tadesse, F. G., Ashine, T., Teka, H., Esayas, E., Messenger, L. A., Chali, W., … Bousema, T. (2021). Anopheles stephensi Mosquitoes as Vectors of Plasmodium vivax and falciparum, Horn of Africa, 2019. Emerg Infect Dis, 27(2), 603–607. doi:10.3201/eid2702.200019

Todd, E. V., Black, M. A., & Gemmell, N. J. (2016). The power and promise of RNA-seq in ecology and evolution. Mol Ecol, 25(6), 1224–1241. doi:10.1111/mec.13526

Villena, O. C., Ryan, S. J., Murdock, C. C., & Johnson, L. R. (2022). Temperature impacts the environmental suitability for malaria transmission by Anopheles gambiae and Anopheles stephensi. Ecology, e3685. doi:10.1002/ecy.3685

WHO. (2021). *World malaria report* 2021 (978-92-4-004049-6). Retrieved from Geneva, Switzerland:

Wickham, H. (2016). ggplot2: Elegant Graphics for Data Analysis. New York: Springer-Verlag.

Wimalasiri-Yapa, B., Barrero, R. A., Stassen, L., Hafner, L. M., McGraw, E. A., Pyke, A. T., … Frentiu, F. D. (2021). Temperature modulates immune gene expression in mosquitoes during arbovirus infection. Open Biol, 11(1), 200246. doi:10.1098/rsob.200246

Yamada, T. G., Hiki, Y., Hiroi, N. F., Shagimardanova, E., Gusev, O., Cornette, R., … Funahashi, A. (2020). Identification of a master transcription factor and a regulatory mechanism for desiccation tolerance in the anhydrobiotic cell line Pv11. PLoS One, 15(3), e0230218. doi:10.1371/journal.pone.0230218

